# GlypNirO: An automated workflow for quantitative *N*- and *O*-linked glycoproteomic data analysis

**DOI:** 10.1101/2020.06.15.153528

**Authors:** Toan K. Phung, Cassandra L. Pegg, Benjamin L. Schulz

## Abstract

Mass spectrometry glycoproteomics is rapidly maturing, allowing unprecedented insights into the diversity and functions of protein glycosylation. However, quantitative glycoproteomics remains challenging. We developed GlypNirO, an automated software pipeline which integrates the complementary outputs of Byonic and Proteome Discoverer to allow high-throughput automated quantitative glycoproteomic data analysis. The output of GlypNirO is clearly structured, allowing manual interrogation, and is also appropriate for input into diverse statistical workflows. We used GlypNirO to analyse a published plasma glycoproteome dataset and identified changes in site-specific *N*- and *O*-glycosylation occupancy and structure associated with hepatocellular carcinoma as putative biomarkers of disease.

## Introduction

Glycosylation is a key post-translational modification critical for protein folding and function in eukaryotes [1-3]. Diverse types of glycosylation are known, all involving modification of specific amino acid residues with complex carbohydrate structures. *N*-linked glycosylation of asparaginies and *O*-linked glycosylation of serines and threonines are the most widely encountered and well studied in eukaryotes. A key feature of glycosylation critical to its biological functions and important for its analysis is its high degree of heterogeneity [4]. This heterogeneity can take the form of variable occupancy, also known as macroheterogeneity – the presence or absence of modification at a particular site in a protein, due to inefficient transfer of the initial glycan structure [5]. In addition, the non-template driven synthesis of glycan structures means that there can be multiple different glycan structures attached at the same site in a pool of mature glycoproteins [6]. This structural heterogeneity is also known as microheterogeneity. This heterogeneity in glycan structure and occupancy can be influenced by many genetic and environmental factors. As such, protein glycosylation is often regulated in response to physiological or pathological conditions [7]. Accurately profiling the site-specific occupancy and structural heterogeneity of glycosylation across glycoproteomes can therefore provide insight into the biology of healthy and diseased states [8].

The current state-of-the art technology for the characterization, identification, and quantification of the glycome or glycoproteome is liquid chromatography coupled to tandem mass spectrometry (LC-MS/MS) [9]. Popular and powerful glycoproteomic workflows typically involve standard proteomic sample preparation and protease digestion, coupled with depletion of abundant proteins or enrichment of glycopeptides to enable their measurement. There have also been several advances in glycopeptide quantification strategies including chemical labelling, label free and data-independent acquisition methods [10]. Progress in MS technology in particular has enabled deep and sensitive measurement of highly complex glycoproteomes, generating large amounts of high quality data [11]. With that comes the need for robust and automated workflows for extracting meaningful results. Numerous software packages have been developed for analysis of outputs from MS technology to automate the process of transformation of raw MS data into ion intensities and matching them with appropriate glycan and peptide sequence databases for glycopeptide identification (reviewed in [12-16]). However, there are few efficient, robust, and automated workflows for glycopeptide quantification. There are several freely available software programs for quantitative label-free glycoproteomics using MS1 or data-dependent acquisition. These include LaCyTools [17], MassyTools [18], and GlycoSpectrumScan [19], which use a predefined list of analytes and masses to interrogate MS1 data, and I-GPA [20], GlycopeptideGraphMS [21], GlycoFragwork [22], and GlycReSoft [23], which integrate identification and abundance/intensity information for glycopeptides (a recent review is provided in [10]). Importantly, the complexity of glycan heterogeneity requires that downstream analysis often involves manual processing in addition to standard informatics workflows.

Here, we developed and used GlypNirO, an automated bioinformatic workflow for label-free quantitative *N*- and *O*-glycoproteomics that focuses on improving robustness and throughput. Our workflow uses a collection of scripts built on an in-house sequence string handling library and the scientific Python data handling package pandas [24], and integrates outputs of two commonly used software packages in glycoproteomic MS data analysis, Proteome Discoverer (Thermo Fisher) and Byonic (Protein Metrics), to extract occupancy and glycoform abundancy of all identified glycopeptides from LC-MS/MS datasets. We applied the workflow to a published dataset comparing the plasma glycoproteomes of liver cancer patients (heptatocellular carcinoma, HCC) and healthy controls [20]. Our analysis revealed differences in occupancy and glycan compositions of several proteins as potential HCC tumor biomarkers.

## Results and Discussion

### GlypNirO

Byonic is powerful software that allows identification of glycopeptides and peptides from complex glyco/proteomic LC-MS/MS datasets but does not perform quantification. Proteome Discoverer allows robust and facile measurement of peptide abundances using MS1 peptide area under the curve (AUC) information. We developed GlypNirO to integrate the outputs from Byonic and Proteome Discoverer to improve the efficiency, ease, and robustness of quantitative glycoproteomic data analysis. GlypNirO takes Byonic and Proteome Discover output files, and user-defined sample information and processing parameters, performs a series of automated data integration and computational steps, and provides informative and intuitive output files with site-or peptide-specific glycoform abundance data. Glyco/peptide identifications from Byonic are linked with AUC data from Proteome Discoverer by matching the experimental scan number. The sites of glycosylation within each peptide assigned by Byonic are identified and clearly labelled. While identification of glycopeptides based on peptide sequence and glycan monosaccharide composition is comparatively reliable with modern LC-MS/MS and data analytics, it is much more difficult to unambiguously assign the precise site of modification within a glycopeptide. GlypNirO therefore provides two options for analysis: site-specific or peptide-specific. If the user trusts Byonic’s site-specific assignment, then all peptide variants that contain that site are included in calculations of its occupancy and glycoform distribution. If the user prefers to perform a peptide-specific analysis, then each proteolytically unique peptide form is treated separately. GlypNirO then calculates the occupancy and proportion of each glycoform at each site/peptide, and provides output files with the protein name, site and/or peptide information, and occupancy and glycoform abundance. Full details are provided in Experimental.

To provide a proof-of-concept use of GlypNirO, we performed an exploratory reanalysis of a previously published dataset [20] obtained from the ProteomeXchange Consortium via the MassIVE repository (PXD003369, MSV000079426). This study performed glycoproteomic LC-MS/MS analysis of whole plasma or plasma depleted of six abundant proteins from liver cancer (hepatocellular carcinoma (HCC)) patients and healthy controls. We identified glycopeptides and peptides in the datafiles from these samples using Byonic, searching separately for *O*- and *N*-glycopeptides (Supplementary Tables S1-S24), and processed the files with Proteome Discoverer (Supplementary Tables S25-S36). We then used GlypNirO to process these results files. This analysis was able to identify and measure 851 *N*-glycopeptides (site-specific) from 150 proteins and 301 *O*-glycopeptides (peptide level) from 89 proteins (Supplementary Tables S37-S40).

Several changes in site-specific glycosylation associated with HCC had been previously identified [20]. We benchmarked the performance of our workflow using GlypNirO with these previously reported changes. Consistent with previous analysis, we found that agalactosylated *N*-glycans on IgG were increased in abundance in HCC (Fig. 1a), and the relative abundance of the HexNAc(5)Hex(6)NeuAc(3) composition at multiple sites on alpha-1-antichymotrypsin was decreased in HCC (Fig. 1b).

**Figure 1.**
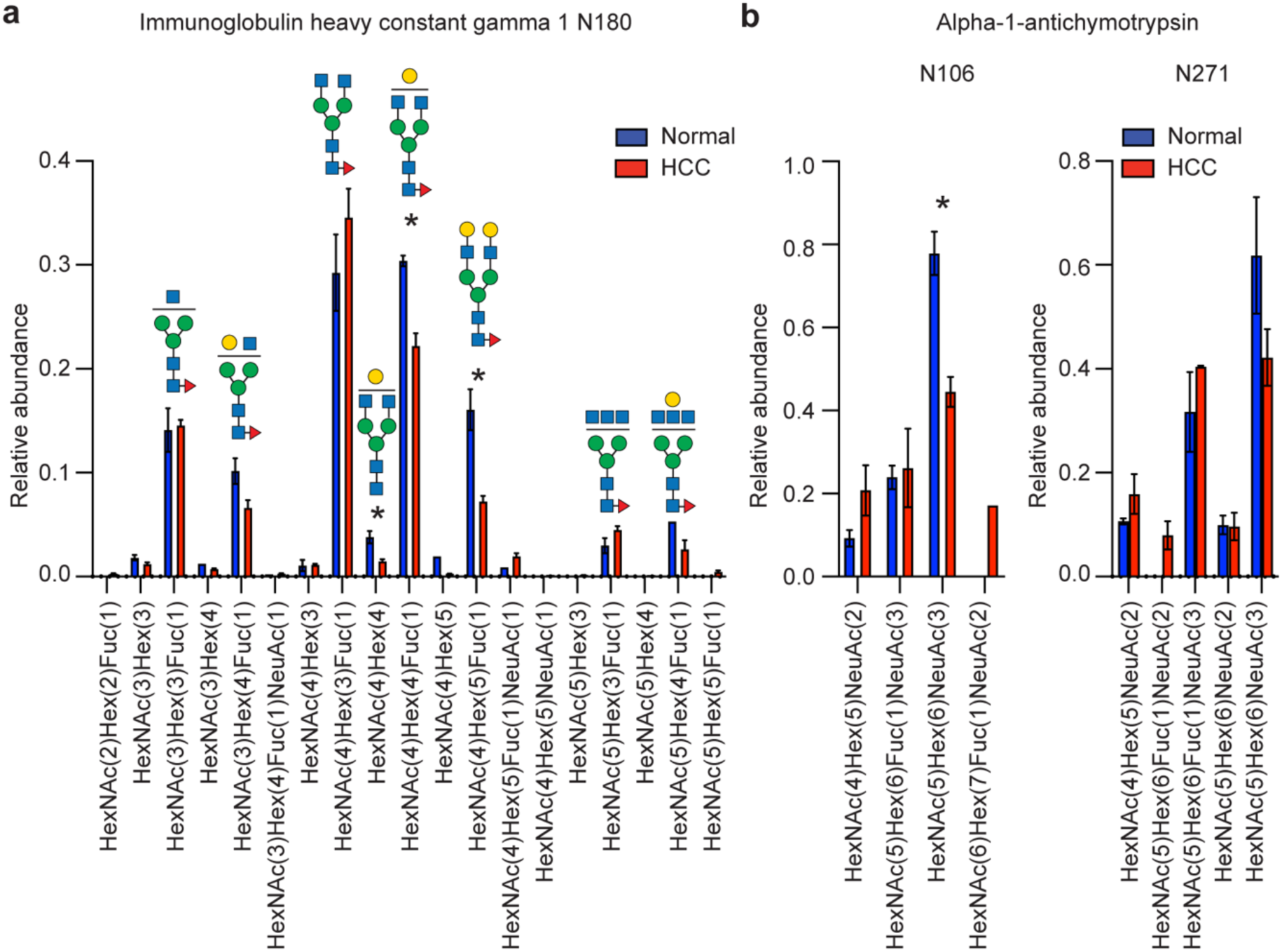
Evaluation of GlypNirO site-specific *N*-glycosylation profiling. Site-specific relative glycoform abundance in HCC and health controls at (**a**) immunoglobulin heavy constant gamma 1 (IgG1) N180, and (**b**) alpha-1-antichymotrypsin N106 and N271.

### *N*-glycoproteome analysis

To extend our analysis, we next investigated the full suite of *N*-glycosylation sites that we were able to identify and measure with GlypNirO. Comparing the site-specific glycoform relative abundance and occupancy, we identified 111 unique glycopeptides with increased and 128 with decreased abundance in HCC compared with healthy controls in depleted plasma, and 93 increased and 67 decreased in HCC in non-depleted plasma (P<0.05, Fig. 2a and b). This analysis with GlypNirO of site-specific relative glycoform abundance confirmed that HCC was associated not only with changes in glycoprotein abundance in plasma, but with changes in the proportions of different glycan structures at specific sites in diverse glycoproteins.

**Figure 2.**
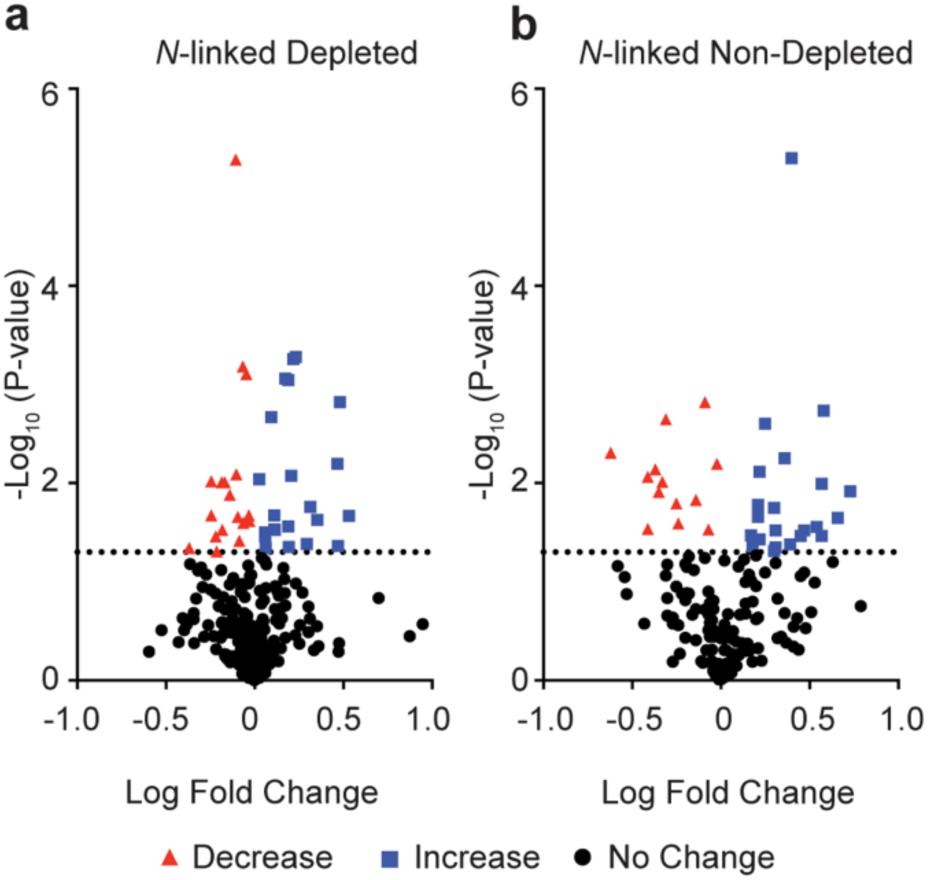
*N*-glycoproteome profiling with GlypNirO. Volcano plots of site-specific *N*-glycoform relative abundance in HCC patients versus healthy controls in (**a**) depleted, and (**b**) non-depleted plasma.

Examining the data in more detail identified several sites with multiple glycoforms with statistically significant changes in abundance. Specifically, HCC patients had decreased abundance of disialylated *N*-glycans at alpha-1-antitrypsin N271 and haptoglobin N184 (Fig. 3a and b), with increased abundance of non-sialylated *N*-glycans at fibrinogen N78 (Fig. 3c), and decreased abundance of trisialylated *N*-glycans at alpha-2-HS-glycoprotein N176 (Fig. 3d). Together, this suggests an overall decrease in sialylation of *N*-glycans across the plasma glycoproteome in HCC.

**Figure 3.**
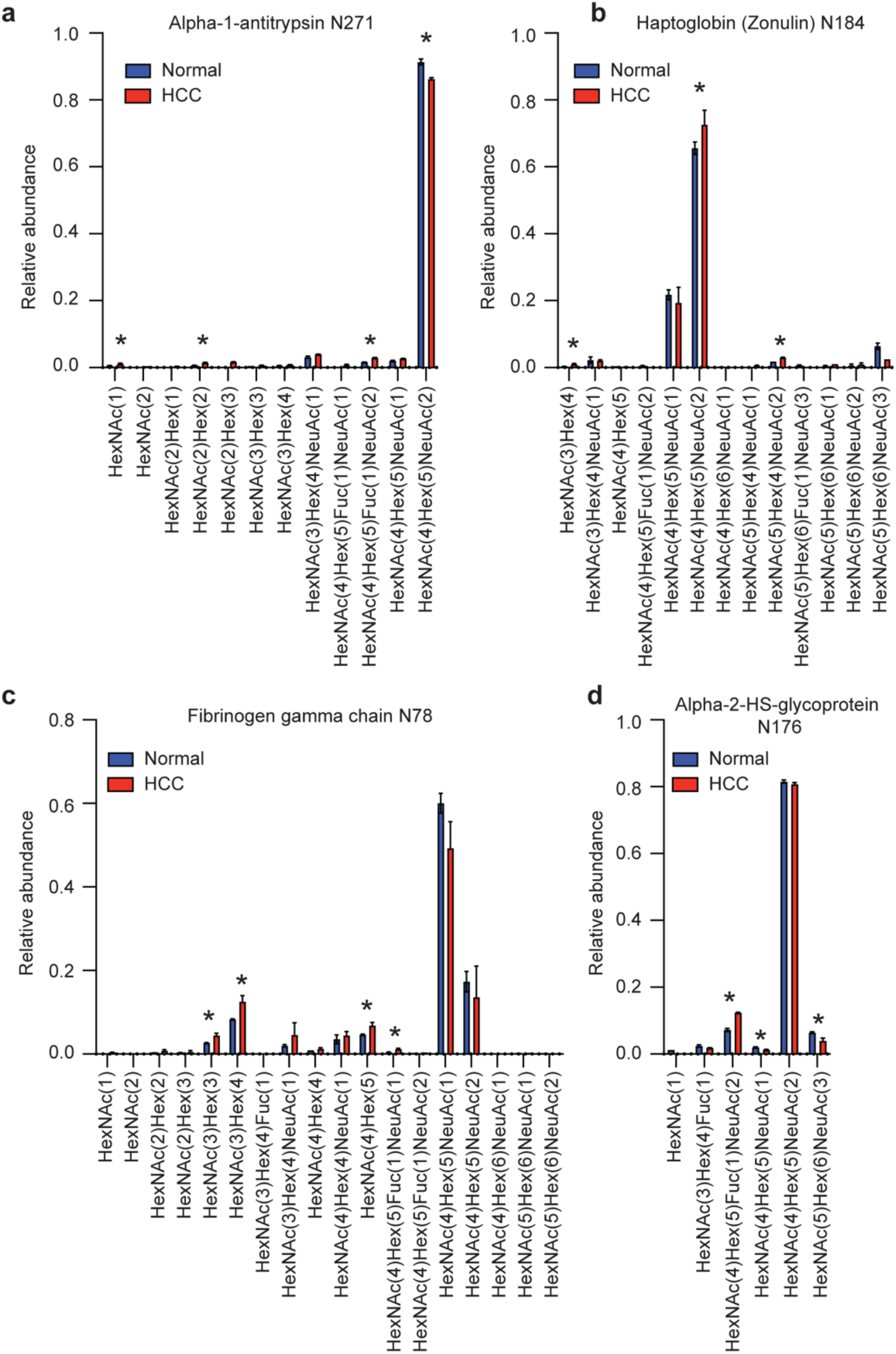
Site-specific *N*-glycopeptide profiling with GlypNirO. Site-specific relative glycoform abundance in HCC patients and health controls at (**a**) alpha-1-antitrypsin N271, (**b**) haptoglobin N184, (**c**) fibrinogen gamma chain N78, and (**d**) alpha-2-HS-glycoprotein N176. N=3; values show mean; error bars show standard error of the mean; *, P<0.05.

### *O*-glycoproteome analysis

The plasma *O*-glycoproteome is perhaps somewhat neglected [25], despite the importance of *O*-glycosylation to diverse aspects of fundamental biology, health, and disease. We therefore investigated all *O*-glycosylation sites that we were able to identify and measure with GlypNirO. Because there are often multiple potential sites of *O*-glycosylation within a tryptic peptide and site-specific assignment is challenging with CID or HCD fragmentation information, we used peptide-centric analysis of the plasma *O*-glycoproteome. Comparing peptide-specific glycoform relative abundance and occupancy, we identified 41 unique *O*-glycopeptides with increased and 27 with decreased abundance in HCC compared with healthy controls in depleted plasma, and 17 increased and 26 decreased in HCC in non-depleted plasma (P<0.05, Fig. 4a and b). As the dataset we analysed measured enriched glycopeptides, it is likely that unglycosylated peptides forms are underrepresented.

**Figure 4.**
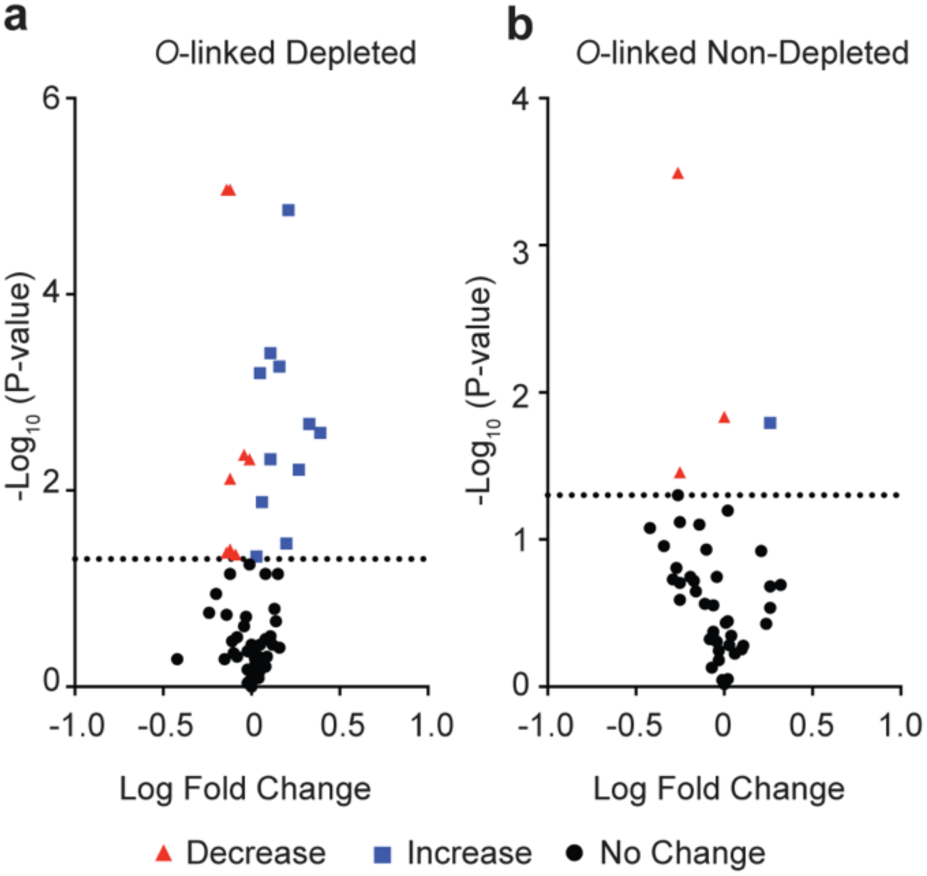
*O*-glycoproteome profiling with GlypNirO. Volcano plots of site-specific *O*-glycoform relative abundance in HCC patients versus healthy controls in (**a**) depleted, and (**b**) non-depleted plasma.

We could identify both changes in peptide-specific *O*-glycan compositions and in *O*-glycan occupancy. HCC patients had increased glycan occupancy and decreased abundance of monosialylated *O*-glycan on fibrinogen alpha chain G_272_GSTSYGTGSETESPR (Fig. 5a). HCC patients showed a relative decrease in disialylated and an increase in monosialylated *O*-glycan abundance on both plasma protease C1 inhibitor V_45_AATVISK and histidine-rich glycoprotein S_271_STTKPPFKPHGSR (Fig. 5b and 5c). Together, and consistent with our *N*-glycoproteome analyses, this suggests that HCC is associated with an overall decrease in sialylation of *N*- and *O*-glycans across the plasma glycoproteome.

**Figure 5.**
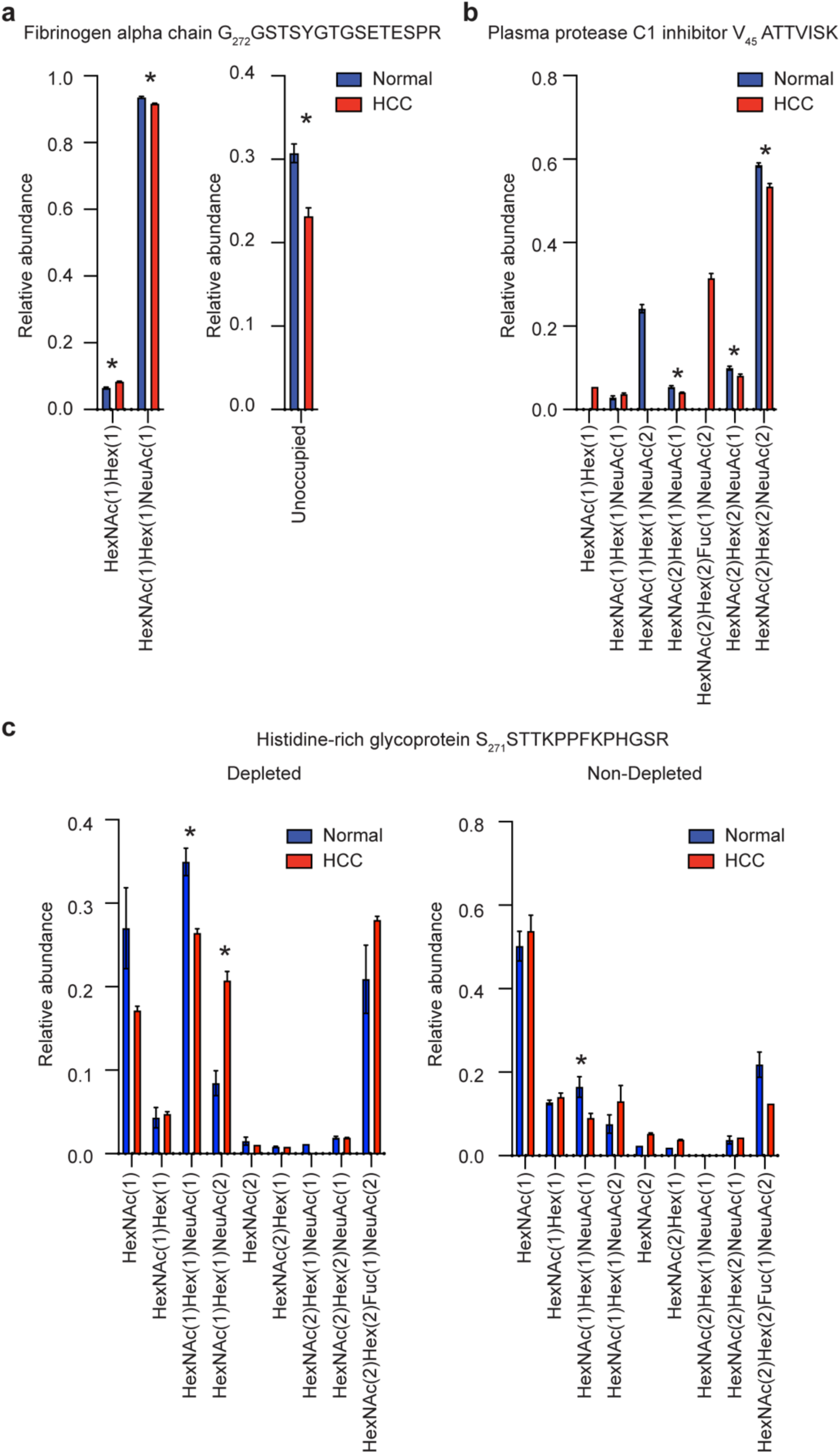
Peptide-specific *O*-glycosylation profiling with GlypNirO. Peptide-specific relative glycoform abundance in HCC patients and health controls on (**a**) fibrinogen alpha chain G_272_GSTSYGTGSETESPR, (**b**) plasma protease C1 inhibitor V_45_AATVISK, and (**c**) histidine-rich glycoprotein S_271_STTKPPFKPHGSR. N=3; values show mean; error bars show standard error of the mean; *, P<0.05.

## Conclusion

GlypNirO is an automated software pipeline that integrates glyco/peptide identification from Byonic and quantification from Proteome Discoverer, and provides output that is appropriate for both manual inspection and further statistical analyses. We note that all glycopeptide identification and quantification workflows will include false positive and negative results, and users should ensure data is appropriately searched and curated before processing with GlypNirO. Additionally, modern LC-MS/MS glycoproteomics cannot fully structurally characterize glycans and often struggles to confidently assign the precise sites of modification; ambiguities which may confound quantification workflows. Our proof-of-principle analysis of a plasma glycoproteome dataset demonstrated that GlypNirO can be used to detect changes in site-specific glycosylation occupancy and structure of *N*- and *O*-glycosylation in complex glycoproteomes. Specifically, we found that HCC was associated with decreased sialylation of both *N*- and *O*-glycans at specific sites on select plasma glycoproteins. GlypNirO will be a useful tool for enabling robust high-throughput quantitative glycoproteomics.

## Experimental

### Byonic and Proteome Discoverer analysis

We identified glycopeptides and peptides using Byonic (Protein Metrics, v. 3.8.13) searching all DDA files (n=12) downloaded from a previously published dataset [20] obtained from the ProteomeXchange Consortium via the MassIVE repository (PXD003369, MSV000079426). Two searches were conducted on each file, one *N*-linked and one *O*-linked. A human protein database was used (UniProt UP000005640, downloaded 20 April 2018 with 20,303 reviewed proteins) [26]. Cleavage specificity was set as C-terminal to Arg/Lys with a maximum of one missed cleavage. The precursor mass tolerance was 10 ppm and fragment ion mass tolerances for CID and HCD were 0.5 Da and 20 ppm, respectively. Cys-S-beta-propionamide was set as a fixed modification, and dynamic modifications included deamidation of asparagine, mono-oxidised methionine, and the formation of pyroglutamate at N-terminal glutamic acid and glutamine. All variable modifications were set as “Common 1” allowing each modification to be present once on a peptide. For *N*-linked searches (N-X-S/T) a database of 164 *N*-glycans was used (Supplementary Table S41) and for the *O*-linked searches (at any S/T) a database of 49 *O*-glycans (Supplementary Table S42) was used. All glycan modifications were set as “Rare 1” allowing each modification to be present once on a peptide. A maximum of two common modifications and one rare modification were allowed per peptide. A protein false discovery rate cut-off of 1% was applied along with the peptide automatic score cutoff [27]. Precursor peak areas were calculated using the Precursor Ions Area Detector node in Proteome Discoverer (v. 2.0.0.802 Thermo Fisher Scientific).Text output files from Proteome Discoverer and Byonic were then used in GlypNirO (https://github.com/bschulzlab/glypniro and Supplementary Information).

### Output combination and preprocessing

GlypNirO was built and used in Python 3.8.3 with backward compatibility tested up to Python 3.6. Each Byonic output file was first iteratively prepared for linking with AUC information from the Proteome Discoverer output. Using a regular expression pattern provided by UniProtKB, the UniProtKB accession ID of each protein from the *Protein Name* column of the Byonic output was parsed and saved into a new temporary *master id* column. If a UniProtKB accession ID could not be matched, the entire protein name was saved into the *master id* column. *Reverse* (decoy) sequences and *Common contaminant proteins* were filtered and removed from the dataset.

To combine data from different isoforms of the same protein, the Byonic output was grouped by accession ID in the *master id* column. From the *Scan number* column, the numeric scan number associated with a PSM was extracted into a temporary *Scan number* column. Area Under the Curve (AUC) information from the *First Scan* column from the Proteome Discoverer output text file was assigned to Byonic data at each corresponding scan number, in the *Area* column. Entries with no AUC value and those not meeting a user-defined Byonic score cutoff (200 here) were removed from the data set.

Using the *Glycans NHFAGNa* and *Modification Type(s)* column, the script obtained the monosaccharide composition of the attached glycan. In the standard Byonic output, only the Δmass of the modification is directly indicated on the modified peptide sequence, with no direct indication of the identity of the corresponding modification. The script therefore calculated the theoretical mass of the glycan from the *Glycans NHFAGNa* column, and matched this to the corresponding amino acid in the peptide. This allowed unambiguous assignment of each site of glycosylation from the Byonic output. Options were provided to either include Byonic assignments of site-specificity, or not, in calculation for the final output.

### Unique PSM selection and glycoform AUC calculation

The compiled dataset as a whole was sorted based on two levels in descending order, first by *Area* and then by *Score*. Two options were available for glyco/peptide grouping: site-specific analysis, or peptide-specific analysis. For site-specific analysis, the site-specificity of glycosylation assigned by Byonic was trusted, and all peptide variants that contained that site were included in calculations of its occupancy and glycoform distribution. PSMs with identical unmodified peptide sequence, glycan monosaccharide composition, calculated *m/z*, and site of glycosylation were grouped. For each group, the PSM precursor *m/z* value with the highest associated *Area* was selected as the unique PSM. The *Area* associated with each unique PSM was used for the calculation of the total AUC of each glycoform at each identified glycosylation site.

For peptide-specific analysis, the precise site of glycosylation within a peptide as assigned by Byonic was ignored, and each proteolytically unique peptide form was treated separately. PSMs with identical unmodified peptide sequence, glycan monosaccharide composition, and calculated *m/z* were grouped. As with site-specific analysis, for each group, the PSM with the highest *Area* was selected as its unique PSM. The *Area* of each unique PSM was used for the calculation of the total AUC of each glycoform for each unique proteolytic peptide.

### Proportional data analysis and final output

In order to allow comparisons of site-specific glycoform abundance and occupancy between different samples, the proportion of each glycoform was calculated with and without inclusion of unglycosylated peptides. For calculation of proportion, glycosylation status was assumed to not quantitatively affect detection. These results were concatenated into the final output file, where columns are the different samples and rows are the different peptide and glycoforms that have been analyzed. The protein name of each glycosylated protein detected in the analysis was also included, parsed from the online UniProtKB database using an inhouse Python library.

### Statistical analyses

Significant differences in glycoform abundances between healthy and diseased samples were evaluated using an unpaired two-tailed t-test without corrections for multiple comparisons. Missing values were not imputed. Spectra were manually validated for glycoforms of interest.

## Supporting information

glypniro-master.zip

## Supporting Information

### Supplementary Tables

Supplementary Table S1 N-link Depleted_Plasma_HCC_1.raw_Byonic.xlsx

Supplementary Table S2 N-link Depleted_Plasma_HCC_2.raw_Byonic.xlsx

Supplementary Table S3 N-link Depleted_Plasma_HCC_3.raw_Byonic.xlsx

Supplementary Table S4 N-link Depleted_Plasma_Nor_1.raw_Byonic.xlsx

Supplementary Table S5 N-link Depleted_Plasma_Nor_2.raw_Byonic.xlsx

Supplementary Table S6 N-link Depleted_Plasma_Nor_3.raw_Byonic.xlsx

Supplementary Table S7 N-link Non_Depleted_Plasma_HCC_1.raw_Byonic.xlsx

Supplementary Table S8 N-link Non_Depleted_Plasma_HCC_2.raw_Byonic.xlsx

Supplementary Table S9 N-link Non_Depleted_Plasma_HCC_3.raw_Byonic.xlsx

Supplementary Table S10 N-link Non_Depleted_Plasma_Nor_1.raw_Byonic.xlsx

Supplementary Table S11 N-link Non_Depleted_Plasma_Nor_2.raw_Byonic.xlsx

Supplementary Table S12 N-link Non_Depleted_Plasma_Nor_3.raw_Byonic.xlsx

Supplementary Table S13 O-link Depleted_Plasma_HCC_1.raw_Byonic.xlsx

Supplementary Table S14 O-link Depleted_Plasma_HCC_2.raw_Byonic.xlsx

Supplementary Table S15 O-link Depleted_Plasma_HCC_3.raw_Byonic.xlsx

Supplementary Table S16 O-link Depleted_Plasma_Nor_1.raw_Byonic.xlsx

Supplementary Table S17 O-link Depleted_Plasma_Nor_2.raw_Byonic.xlsx

Supplementary Table S18 O-link Depleted_Plasma_Nor_3.raw_Byonic.xlsx

Supplementary Table S19 O-link Non_Depleted_Plasma_HCC_1.raw_Byonic.xlsx

Supplementary Table S20 O-link Non_Depleted_Plasma_HCC_2.raw_Byonic.xlsx

Supplementary Table S21 O-link Non_Depleted_Plasma_HCC_3.raw_Byonic.xlsx

Supplementary Table S22 O-link Non_Depleted_Plasma_Nor_1.raw_Byonic.xlsx

Supplementary Table S23 O-link Non_Depleted_Plasma_Nor_2.raw_Byonic.xlsx

Supplementary Table S24 O-link Non_Depleted_Plasma_Nor_3.raw_Byonic.xlsx

Supplementary Table S25 Depleted_Plasma_HCC_1_MSnSpectrumInfo.txt

Supplementary Table S26 Depleted_Plasma_HCC_2_MSnSpectrumInfo.txt

Supplementary Table S27 Depleted_Plasma_HCC_3_MSnSpectrumInfo.txt

Supplementary Table S28 Depleted_Plasma_Nor_1_MSnSpectrumInfo.txt

Supplementary Table S29 Depleted_Plasma_Nor_2_MSnSpectrumInfo.txt

Supplementary Table S30 Depleted_Plasma_Nor_3_MSnSpectrumInfo.txt

Supplementary Table S31 Non_Depleted_Plasma_HCC_1_MSnSpectrumInfo.txt

Supplementary Table S32 Non_Depleted_Plasma_HCC_2_MSnSpectrumInfo.txt

Supplementary Table S33 Non_Depleted_Plasma_HCC_3_MSnSpectrumInfo.txt

Supplementary Table S34 Non_Depleted_Plasma_Nor_1_MSnSpectrumInfo.txt

Supplementary Table S35 Non_Depleted_Plasma_Nor_2_MSnSpectrumInfo.txt

Supplementary Table S36 Non_Depleted_Plasma_Nor_3_MSnSpectrumInfo.txt

Supplementary Table S37 N-linked site-specific output.xlsx

Supplementary Table S38 O-linked peptide output.xlsx

Supplementary Table S39 O-linked site-specific output.xlsx

Supplementary Table S40 N-linked peptide output.

Supplementary Table S41 N-linked Glycan Database.xlsx

Supplementary Table S42 O-linked Glycan Database.xlsx

### Supplementary Figures

**Supplementary Figure S1.**
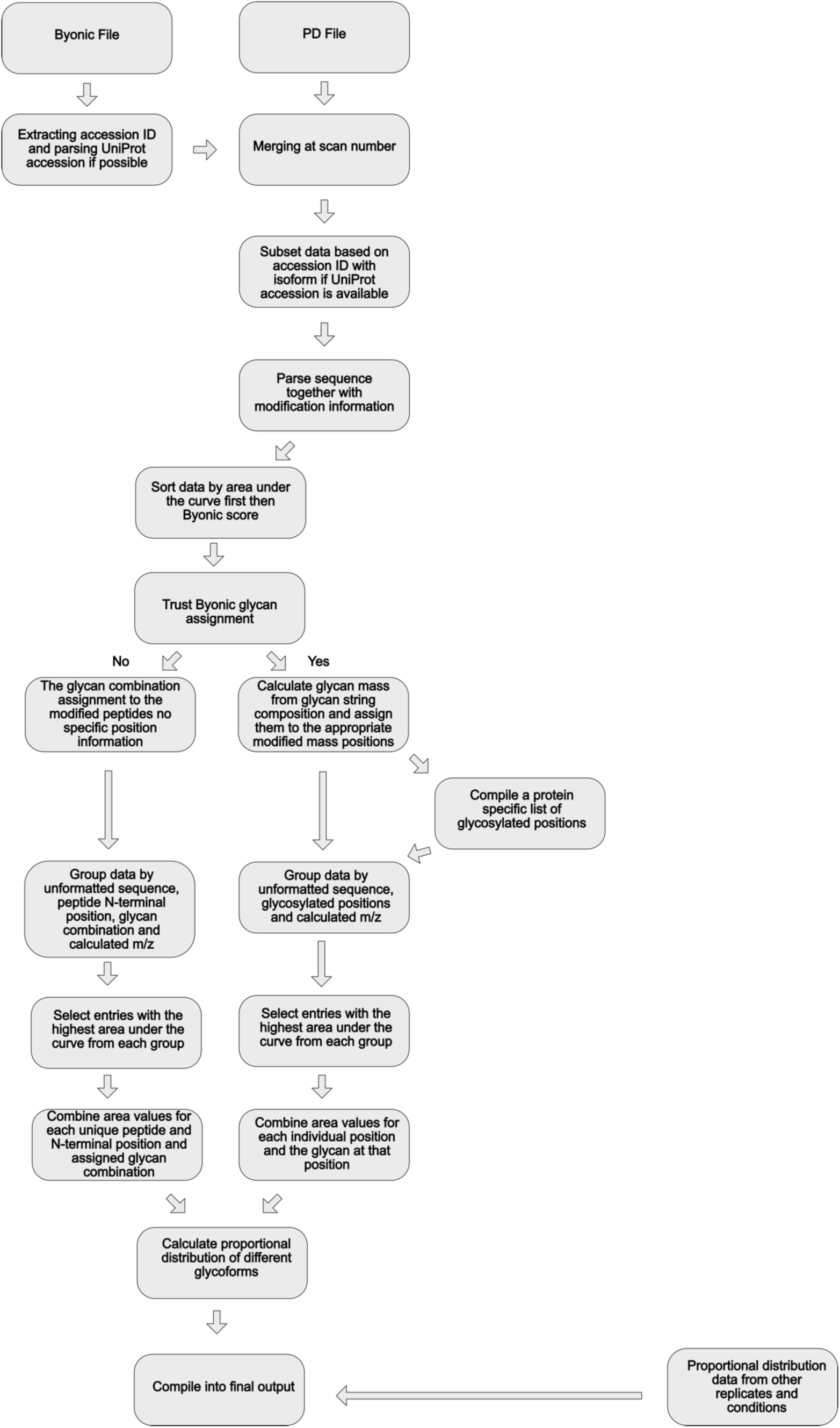
GlypNirO workflow overview.

**Supplementary Information**

glypniro-master.zip

