## Supplementary material for "GlypNirO: An automated workflow for quantitative *N*- and *O*-linked glycoproteomic data analysis": glypniro-master.zip: common.py

```
from io import StringIO
from copy import deepcopy
import numpy as np
import pandas as pd
import re

from glypnirO_GUI.get_uniprot import UniprotParser
from sequal.sequence import Sequence
from sequal.resources import glycan_block_dict

# Defining important colume names within the dataset
sequence_column_name = "Peptide\n< ProteinMetrics Confidential >"
glycans_column_name = "Glycans\nNHFAGNa"
starting_position_column_name = "Starting\nposition"
modifications_column_name = "Modification Type(s)"
observed_mz = "Calc.\nmass (M+H)"
protein_column_name = "Protein Name"
rt = "Scan Time"
selected_aa = {"N", "S", "T"}

# Defining important regular expression pattern to parse the dataset
regex_glycan_number_pattern = "\d+"
glycan_number_regex = re.compile(regex_glycan_number_pattern)
regex_pattern = "\.[\[\]\w\.\+\-]*\."
sequence_regex = re.compile(regex_pattern)
uniprot_regex = re.compile("(?P<accession>[OPQ][0-9][A-Z0-9]{3}[0-9]|[A-NR-Z][0-9]([A-Z][A-Z0-9]{2}[0-9]){1,2})(?P<isoform>-\d)?")
glycan_regex = re.compile("(\w+)\((\d+)\)")


# Function to filter for only PSM collections that do not only containing unglycosylated peptides
def filter_U_only(df):
    unique_glycan = df["Glycans"].unique()
    if len(unique_glycan) > 1 or True not in np.isin(unique_glycan, "U"):
        # print(unique_glycan)
        return True
    return False

# Filter for only PSMs that are unglycosylated within PSM collections that do not only containing unglycosylated peptides
def filter_with_U(df):
    unique_glycan = df["Glycans"].unique()
    if len(unique_glycan) > 1 \
            and \
            True in np.isin(unique_glycan, "U"):
        return True
    return False

# parse modification mass and convert it from string to float
def get_mod_value(amino_acid):
    if amino_acid.mods:
        if amino_acid.mods[0].value.startswith("+"):
            return float(amino_acid.mods[0].value[1:])
        else:
            return -float(amino_acid.mods[0].value[1:])
    else:
        return 0

# load fasta file into a dictionary
def load_fasta(fasta_file_path, selected=None, selected_prefix=""):
    with open(fasta_file_path, "rt") as fasta_file:
        result = {}
        current_seq = ""
        for line in fasta_file:
            line = line.strip()

            if line.startswith(">"):
                if selected:
                    if selected_prefix + line[1:] in selected:
                        result[line[1:]] = ""
                        current_seq = line[1:]
                else:
                    result[line[1:]] = ""
                    current_seq = line[1:]
            else:
                result[current_seq] += line
        return result

# Storing analysis result for each protein
class Result:
    def __init__(self, df):
        self.df = df
        self.empty = df.empty

    def calculate_proportion(self, occupancy=True):
        """
        calculate proportion of each glycoform from the dataset
        :type occupancy: bool
        whether or not to calculate the proportion as occupancy which would includes unglycosylated form.

        """
        df = self.df.copy()
        #print(df)
        if not occupancy:
            df = df[df["Glycans"] != "U"]
        if "Peptides" in df.columns:
            gr = [# "Isoform",
                  "Peptides", "Position"]
        else:
            gr = [# "Isoform",
                  "Position"]
        for _, g in df.groupby(gr):
            total = g["Value"].sum()
            for i, r in g.iterrows():
                df.at[i, "Value"] = r["Value"] / total

        return df

    def to_summary(self, df=None, name="", trust_byonic=False, occupancy=True):
        """

        :type trust_byonic: bool
        whether or not to calculate calculate raw values for each individual position assigned by byonic
        :type occupancy: bool
        whether or not to calculate the proportion as occupancy which would includes unglycosylated form.
        :type df: pd.DataFrame

        """
        if df is None:
            df = self.df
        if not occupancy:
            df = df[df["Glycans"] != "U"]
        if trust_byonic:
            temp = df.set_index([# "Isoform",
                                 "Position", "Glycans"])
        else:
            temp = df.set_index([# "Isoform",
                                 "Peptides", "Glycans", "Position"])

        temp.rename(columns={"Value": name}, inplace=True)
        return temp

# Object containing each individual protein. much of the methods involved in the analysis is contained within this object
# Each protein is assigned one of the GlypnirOcomponent object with a subset of their PD and Byonic data
class GlypnirOComponent:
    def __init__(self, filename, area_filename, replicate_id, condition_id, protein_name, minimum_score=0, trust_byonic=False, legacy=False):
        if type(filename) == pd.DataFrame:
            data = filename.copy()
        else:
            data = pd.read_excel(filename, sheet_name="Spectra")
        if type(area_filename) == pd.DataFrame:
            file_with_area = area_filename
        else:
            if area_filename.endswith("xlsx"):
                file_with_area = pd.read_excel(area_filename)
            else:
                file_with_area = pd.read_csv(area_filename, sep="\t")

        # Joining of area and glycan data for each PSM using scan number as merging point
        data["Scan number"] = pd.to_numeric(data["Scan #"].str.extract("scan=(\d+)", expand=False))
        data = pd.merge(data, file_with_area, left_on="Scan number", right_on="First Scan")

        # Subset and filtering data for those with non blank area value and passing minimum score cutoff
        self.protein_name = protein_name
        self.data = data.sort_values(by=['Area'], ascending=False)
        self.replicate_id = replicate_id
        self.condition_id = condition_id
        self.data = data[data["Area"].notnull()]
        self.data = self.data[(self.data["Score"] >= minimum_score) &
                         (self.data[protein_column_name].str.contains(protein_name))
                         # (data["Protein Name"] == ">"+protein_name) &
                         ]
        self.data = self.data[~self.data[protein_column_name].str.contains(">Reverse")]
        if len(self.data.index) > 0:
            self.empty = False
        else:
            self.empty = True
        self.row_to_glycans = {}
        self.glycan_to_row = {}

        self.trust_byonic = trust_byonic
        self.legacy = legacy
        self.sequon_glycosites = set()
        self.glycosylated_seq = set()

    # method for calculate glycan mass from string syntax using regular expression and a dictionary of glycan block and their mass
    def calculate_glycan(self, glycan):
        current_mass = 0
        current_string = ""
        for i in glycan:
            current_string += i
            if i == ")":
                s = glycan_regex.search(current_string)
                if s:
                    name = s.group(1)
                    amount = s.group(2)
                    current_mass += glycan_block_dict[name]*int(amount)
                current_string = ""
        return current_mass

    # process the protein data
    def process(self):

        for i, r in self.data.iterrows():
            glycan_dict = {}
            search = sequence_regex.search(r[sequence_column_name])
            # get peptide sequence without flanking prefix and suffix amino acids then create a Sequence object from the string
            seq = Sequence(search.group(0))
            # get unformated string from the Sequence object. This unformatted string contain a "." at both end
            stripped_seq = seq.to_stripped_string()

            # Get parse glycans from glycan column into a list
            glycans = []
            if pd.notnull(r[glycans_column_name]):
                glycans = r[glycans_column_name].split(",")
            if search:
                # store the unformatted sequence without "." at both end into the dataframe
                self.data.at[i, "stripped_seq"] = stripped_seq.rstrip(".").lstrip(".")
                # calculate the programmatic starting position of the sequence
                origin_seq = r[starting_position_column_name] - 1
                glycan_reordered = []
                self.data.at[i, "origin_start"] = origin_seq
                #calculate the programmatic stopping position of the sequence
                self.data.at[i, "Ending Position"] = r[starting_position_column_name] + len(self.data.at[i, "stripped_seq"])
                self.data.at[i, "position_to_glycan"] = ""
                if self.trust_byonic:
                    n_site_status = {}
                    p_n = r[protein_column_name].lstrip(">")

                    # current_glycan = 0
                    max_glycans = len(glycans)
                    glycosylation_count = 1
                    # creating dictionary storing the glycan and its mass
                    if max_glycans:
                        self.row_to_glycans[i] = np.sort(glycans)
                        for g in glycans:
                            data_gly = self.calculate_glycan(g)
                            glycan_dict[str(round(data_gly, 3))] = g
                            self.glycan_to_row[g] = i

                    glycosylated_site = []
                    # iterating through the unformated sequence and assign glycan to modified position based on the modified mass
                    for aa in range(1, len(seq) - 1):
                        if seq[aa].mods:
                            mod_value = float(seq[aa].mods[0].value)
                            round_mod_value = round(mod_value)

                            round_3 = round(mod_value, 3)
                            # if the glycan is identified to be found, store the position of the glycosylated amino acid on the protein sequence for later reference
                            if str(round_3) in glycan_dict:
                                seq[aa].extra = "Glycosylated"
                                pos = int(r[starting_position_column_name]) + aa - 2
                                self.sequon_glycosites.add(pos + 1)
                                position = "{}_position".format(str(glycosylation_count))
                                self.data.at[i, position] = seq[aa].value + str(pos + 1)
                                glycosylated_site.append(self.data.at[i, position] + "_" + str(round_mod_value))
                                glycosylation_count += 1
                                glycan_reordered.append(glycan_dict[str(round_3)])
                    if glycan_reordered:
                        self.data.at[i, "position_to_glycan"] = ",".join(glycan_reordered)
                    self.data.at[i, "glycoprofile"] = ";".join(glycosylated_site)

                else:
                    # if the analysis is only done on peptide and glycan combination, we would only need to set whether the peptide is glycosylated and store the unformatted peptide sequence of the glycosylated one for later reference
                    if pd.notnull(r[glycans_column_name]):
                        glycans = r[glycans_column_name].split(",")
                        glycans.sort()
                        self.data.at[i, glycans_column_name] = ",".join(glycans)
                        self.data.at[i, "glycosylation_status"] = True
                        self.glycosylated_seq.add(self.data.at[i, "stripped_seq"])

    # analyze the compiled data by identifying unique PSM and calculate cumulative raw area under the curve
    def analyze(self, max_sites=0, combine_d_u=True, splitting_sites=False):
        result = []
        # sort the data first by area then score in descending order.
        temp = self.data.sort_values(["Area", "Score"], ascending=False)
        # cells in glycan column with no glycan will be assigned a string "None"
        temp[glycans_column_name] = temp[glycans_column_name].fillna("None")
        out = []

        if self.trust_byonic:
            # if trust byonic we would analyze by grouping the data at unformatted sequence, glycosylated positions and calculated m/z
            seq_glycosites = list(self.sequon_glycosites)
            seq_glycosites.sort()

            for i, g in temp.groupby(["stripped_seq", "glycoprofile", observed_mz]):
                seq_within = []
                # select row with highest area value in a group
                unique_row = g.loc[g["Area"].idxmax()]

                if seq_glycosites:
                    for n in seq_glycosites:
                        # create a list of n glycosylation sites that can be found within the peptide sequence
                        if unique_row[starting_position_column_name] <= n < unique_row["Ending Position"]:
                            # print(unique_row["stripped_seq"], n, unique_row[starting_position_column_name])
                            seq_within.append(
                                unique_row["stripped_seq"][n-unique_row[starting_position_column_name]]+str(n))

                glycosylation_count = 0
                glycans = unique_row["position_to_glycan"].split(",")
                # create a dataset of position, glycans associate to that position and area under the curve of them
                for c in range(len(unique_row.index)):
                    if unique_row.index[c].endswith("_position"):
                        if pd.notnull(unique_row[unique_row.index[c]]):
                            pos = unique_row[unique_row.index[c]]
                            result.append({"Position": pos, "Glycans": glycans[glycosylation_count], "Value": unique_row["Area"]})
                            ind = seq_within.index(pos)
                            seq_within.pop(ind)
                            glycosylation_count += 1

                if seq_within:
                    for s in seq_within:
                        result.append({"Position": s, "Glycans": "U", "Value": unique_row["Area"]})

            if result:
                result = pd.DataFrame(result)
                # sum area under the curve of those with the same glycosylation position and glycan composition
                group = result.groupby(["Position", "Glycans"])

                out = group.agg(np.sum).reset_index()
            else:
                out = pd.DataFrame([], columns=["Position", "Glycans", "Values"])

        else:

            # if a peptide level analysis was done, the grouping would be on unformatted sequence, glycan combination, position of the peptide N-terminus, calculated m/z
            for i, g in temp.groupby(["stripped_seq", glycans_column_name, starting_position_column_name, observed_mz]):
                # select and create a dataset of unique psm compositing of the unformatted sequence, glycans, area under the curve and position of the peptide N-terminus
                unique_row = g.loc[g["Area"].idxmax()]

                if unique_row[glycans_column_name] != "None":
                    result.append({"Peptides": i[0], "Glycans": i[2], "Value": unique_row["Area"], "Position": i[3]})
                else:
                    result.append({"Peptides": i[0], "Glycans": "U", "Value": unique_row["Area"], "Position": i[3]})

            result = pd.DataFrame(result)
            # sum those area under the curve with the same peptides, position and glycans
            group = result.groupby(["Peptides", "Position", "Glycans"])
            out = group.agg(np.sum).reset_index()

        return Result(out)


class GlypnirO:
    def __init__(self, trust_byonic=False, get_uniprot=False):
        self.trust_byonic = trust_byonic
        self.components = None
        self.uniprot_parsed_data = pd.DataFrame([])
        self.get_uniprot = get_uniprot

    def add_component(self, filename, area_filename, replicate_id, sample_id):
        component = GlypnirOComponent(filename, area_filename, replicate_id, sample_id)

    # loading of input experiment file
    def add_batch_component(self, component_list, minimum_score, protein=None, combine_uniprot_isoform=True, legacy=False):
        self.load_dataframe(component_list)
        protein_list = []
        if protein is not None:
            self.components["Protein"] = pd.Series([protein]*len(self.components.index), index=self.components.index)
            for i, r in self.components.iterrows():
                comp = GlypnirOComponent(r["filename"], r["area_filename"], r["replicate_id"], condition_id=r["condition_id"], protein_name=protein, minimum_score=minimum_score, trust_byonic=self.trust_byonic, legacy=legacy)
                self.components.at[i, "component"] = comp
                print("{} - {}, {} peptides has been successfully loaded".format(r["condition_id"], r["replicate_id"], str(len(comp.data.index))))

        else:
            components = []
            for i, r in self.components.iterrows():
                data = pd.read_excel(r["filename"], sheet_name="Spectra")
                protein_id_column = protein_column_name
                if combine_uniprot_isoform:
                    protein_id_column = "master_id"
                    for i2, r2 in data.iterrows():
                        # search for uniprot accession id in protein column name
                        search = uniprot_regex.search(r2[protein_column_name])
                        # if the protein is not a decoy or labelled as common contaminant, the accession id would be saved into a master_id column. If no accession id, the whole protein name would be saved there instead.
                        if not r2[protein_column_name].startswith(">Reverse") and not r2[protein_column_name].endswith("(Common contaminant protein)"):
                            if search:
                                data.at[i2, "master_id"] = search.groupdict(default="")["accession"]
                                if not self.get_uniprot:
                                    protein_list.append([search.groupdict(default="")["accession"], r2[protein_column_name]])
                                if search.groupdict(default="")["isoform"] != "":
                                    data.at[i2, "isoform"] = int(search.groupdict(default="")["isoform"][1:])
                                else:
                                    data.at[i2, "isoform"] = 1

                            else:
                                data.at[i2, "master_id"] = r2[protein_column_name]
                                data.at[i2, "isoform"] = 1
                        else:
                            data.at[i2, "master_id"] = r2[protein_column_name]
                            data.at[i2, "isoform"] = 1

                # read pd file
                if r["area_filename"].endswith("xlsx"):
                    file_with_area = pd.read_excel(r["area_filename"])
                else:
                    file_with_area = pd.read_csv(r["area_filename"], sep="\t")

                for index, g in data.groupby([protein_id_column]):

                    u = index
                    if not u.startswith(">Reverse") and not u.endswith("(Common contaminant protein)"):
                        # merging of byonic and pd data for appropriate protein
                        comp = GlypnirOComponent(g, file_with_area, r["replicate_id"],
                                                 condition_id=r["condition_id"], protein_name=u,
                                                 minimum_score=minimum_score, trust_byonic=self.trust_byonic, legacy=legacy)
                        if not comp.empty:
                            components.append({"filename": r["filename"], "area_filename": r["area_filename"], "condition_id": r["condition_id"], "replicate_id": r["replicate_id"], "Protein": u, "component": comp})
                yield i, r
                print(
                    "{} - {} peptides has been successfully loaded".format(r["condition_id"],
                                                                                      r["replicate_id"]))
            self.components = pd.DataFrame(components, columns=list(self.components.columns) + ["component", "Protein"])
            if not self.get_uniprot:
                protein_df = pd.DataFrame(protein_list, columns=["Entry", "Protein names"])
                self.uniprot_parsed_data = protein_df
                #print(self.uniprot_parsed_data)

    def load_dataframe(self, component_list):
        if type(component_list) == list:
            self.components = pd.DataFrame(component_list)
        elif type(component_list) == pd.DataFrame:
            self.components = component_list
        elif type(component_list) == str:
            if component_list.endswith(".txt"):
                self.components = pd.read_csv(component_list, sep="\t")
            elif component_list.endswith(".csv"):
                self.components = pd.read_csv(component_list)
            elif component_list.endswith(".xlsx"):
                self.components = pd.read_excel(component_list)
            else:
                raise ValueError("Input have to be list, pandas dataframe, or csv, xlsx, or tabulated txt filepath.")
        else:
            raise ValueError("Input have to be list, pandas dataframe, or csv, xlsx, or tabulated txt filepath.")

    def process_components(self):
        for i, r in self.components.iterrows():
            # print("Processing {} - {} {} for {}".format(r["condition_id"], r["replicate_id"], r["Protein"], analysis))
            r["component"].process()

    # analysis of the compiled data
    def analyze_components(self):

        result = []
        result_without_u = []
        result_occupancy_no_calculation_u = []
        for i, r in self.components.iterrows():
            print("Analyzing", r["Protein"], r["condition_id"], r["replicate_id"], r["component"].protein_name)
            analysis_result = r["component"].analyze()
            if not analysis_result.empty:
                # get raw and proportional calculation of unique psm auc with unglycosylated peptide
                a = analysis_result.to_summary(name="Raw", trust_byonic=self.trust_byonic)
                pro = analysis_result.calculate_proportion()
                b = analysis_result.to_summary(pro, "Proportion", trust_byonic=self.trust_byonic)
                temp_df = self._summary(a, r, b)
                result.append(temp_df)

                # proportion for glycoforms here are calculated without unglycosylated form.
                a_without_u = analysis_result.to_summary(name="Raw", trust_byonic=self.trust_byonic, occupancy=False)
                pro_without_u = analysis_result.calculate_proportion(occupancy=False)
                b_without_u = analysis_result.to_summary(pro_without_u, "Proportion", trust_byonic=self.trust_byonic, occupancy=False)
                temp_df_without_u = self._summary(a_without_u, r, b_without_u)
                result_without_u.append(temp_df_without_u)

                temp_df_no_calculation_u = self._summary(a, r, b_without_u)
                result_occupancy_no_calculation_u.append(temp_df_no_calculation_u)

        result_occupancy = self._summary_format(result)
        result_occupancy_with_u = self._summary_format(result, filter_with_U, True)
        result_glycoform = self._summary_format(result_without_u)

        tempdf_index_reset_result_occupancy_with_u = result_occupancy_with_u.reset_index()
        tempdf_index_reset_result_glycoform = result_glycoform.reset_index()
        result_occupancy_glycoform_sep = pd.concat(
            [tempdf_index_reset_result_glycoform, tempdf_index_reset_result_occupancy_with_u])
        # format the output with the correct column name for site specific or peptide level analysis
        if self.trust_byonic:
            result_occupancy_glycoform_sep = result_occupancy_glycoform_sep.set_index(["Protein", "Protein names",
                                                                                       # "Isoform",
                                                                                       "Glycosylated positions in peptide", "Glycans"])
            result_occupancy_glycoform_sep = result_occupancy_glycoform_sep.sort_index(
                level=["Protein", "Protein names",
                       # "Isoform",
                       "Glycosylated positions in peptide"])
        else:

            result_occupancy_glycoform_sep = result_occupancy_glycoform_sep.set_index(
                ["Protein", "Protein names",
                 # "Isoform",
                 "Position peptide N-terminus", "Peptides", "Glycans"])
            result_occupancy_glycoform_sep = result_occupancy_glycoform_sep.sort_index(
                level=["Protein", "Protein names",
                       # "Isoform",
                       "Position peptide N-terminus", "Peptides"])


        print("Finished analysis.")
        return {"Glycoforms":
                    result_glycoform,
                "Occupancy":
                    result_occupancy,
                "Occupancy_With_U":
                    result_occupancy_with_u,
                "Occupancy_Without_Proportion_U":
                    result_occupancy_glycoform_sep}

    # summarize the data and collect uniprot protein directly fromt the online uniprot database if get_uniprot is True
    def _summary_format(self, result, filter_method=filter_U_only, select_for_u=False):
        result_data = pd.concat(result)
        result_data = result_data.reset_index(drop=True)
        accessions = result_data["Protein"].unique()

        if self.uniprot_parsed_data.empty:
            if self.get_uniprot:
                parser = UniprotParser(accessions, True)

                data = []
                for i in parser.parse("tab"):
                    frame = pd.read_csv(StringIO(i), sep="\t")
                    frame = frame.rename(columns={frame.columns[-1]: "query"})
                    data.append(frame)
                self.uniprot_parsed_data = pd.concat(data, ignore_index=True)
                #
                self.uniprot_parsed_data = self.uniprot_parsed_data[['Entry', 'Protein names']]
        else:
            self.uniprot_parsed_data = self.uniprot_parsed_data.groupby(["Entry"]).head(1).reset_index().drop(["index"], axis=1)

        result_data = result_data.merge(self.uniprot_parsed_data, left_on="Protein", right_on="Entry")
        result_data.drop("Entry", 1, inplace=True)

        if self.trust_byonic:
            groups = result_data.groupby(by=["Protein", "Protein names",
                                             # "Isoform",
                                             "Position"])
        else:
            groups = result_data.groupby(by=["Protein", "Protein names",
                                             # "Isoform",
                                             "Position", "Peptides"])


        result_data = groups.filter(filter_method)
        if select_for_u:
            result_data = result_data[result_data["Glycans"] == "U"]
        if self.trust_byonic:
            result_data = result_data.rename({"Position": "Glycosylated positions in peptide"}, axis="columns")
            result_data = result_data.set_index(
                ["Label", "condition_id", "replicate_id", "Protein", "Protein names",
                 # "Isoform",
                 "Glycosylated positions in peptide", "Glycans"])
        else:
            result_data = result_data.rename({"Position": "Position peptide N-terminus"}, axis="columns")
            result_data = result_data.set_index(
                ["Label", "condition_id", "replicate_id", "Protein", "Protein names",
                 # "Isoform",
                 "Position peptide N-terminus", "Peptides", "Glycans"])
        result_data = result_data.unstack(["Label", "condition_id", "replicate_id"])
        result_data = result_data.sort_index(level=["Label", "condition_id", "replicate_id"], axis=1)
        #result_data.to_csv("test.txt", sep="\t")
        if self.trust_byonic:
            result_data = result_data.sort_index(level=["Protein", "Protein names",
                                                        # "Isoform",
                                                        "Glycosylated positions in peptide"])
        else:
            result_data = result_data.sort_index(level=["Protein", "Protein names",
                                                        # "Isoform",
                                                        "Position peptide N-terminus", "Peptides"])

        result_data.columns = result_data.columns.droplevel()
        return result_data

    # combine output from different protein, condition and replicate 
    def _summary(self, a, r, b):
        temp_df = pd.concat([a, b], axis=1)

        temp_df.columns.name = "Label"
        temp_df = temp_df.stack()
        lc = [temp_df]
        for c in ["Protein", "condition_id", "replicate_id"]:
            lc.append(pd.Series([r[c]] * len(temp_df.index), index=temp_df.index, name=c))
        temp_df = pd.concat(lc, axis=1)

        temp_df = temp_df.reset_index()

        return temp_df
```
