## Supplementary material for "GlypNirO: An automated workflow for quantitative *N*- and *O*-linked glycoproteomic data analysis": glypniro-master.zip: get_uniprot.py

```
import requests
import re


# regex pattern for matching UniProt accession that can be used with the search object groupdict method to retrieve accession and isotype information separately
acc_regex = re.compile("(?P<accession>[OPQ][0-9][A-Z0-9]{3}[0-9]|[A-NR-Z][0-9]([A-Z][A-Z0-9]{2}[0-9]){1,2})(?P<isotype>-\d)?")

# sequence object for storing and presenting uniprot id.
class UniprotSequence:
    def __init__(self, acc, parse_acc=False):
        """

        :type parse_acc: bool
        whether or not for the script to parse the accession id from the input
        :type acc: str
        a string containing the Uniprot accession ID of the sequence
        """
        self.raw_acc = acc
        self.accession = None
        self.isotype = None
        if parse_acc:
            match = acc_regex.search(self.raw_acc)
            if match:
                self.accession = match.groupdict(default="")["accession"]
                self.isotype = match.groupdict(default="")["isotype"]

    def __str__(self):
        return self.accession + self.isotype

    def __repr__(self):
        return self.accession + self.isotype


# The main operating object for parsing Uniprot ID from the output.
# UniprotParser work by accessing the Uniprot database online api at the base url https://www.uniprot.org/uploadlists/
# The parser divide the input list of accession ids into several smaller lists of max 300 accession ids.
class UniprotParser:
    base_url = "https://www.uniprot.org/uploadlists/"

    def __init__(self, acc_list, unique=False, headers=None):
        """


        :type headers: dict
        headers dictionary
        :type unique: bool
        specify whether the list is unique
        :type acc_list: list
        list of UniProt accession id
        """
        if not headers:
            self.headers = {"User-Agent": "Python, GlypnirO"}
        else:
            self.headers = headers
        self.acc_list = acc_list
        if not unique:
            self.acc_list = list(set(i for i in self.acc_list))
        self.total_input = len(acc_list)

    @staticmethod
    def create_params(acc_list, format="tab", include_isoform=True):
        """

        :type acc_list: list
        list of accession ids
        :type include_isoform: bool
        whether to include isoform or not. Only important in fasta output.
        :type format: str
        the format of the output file

        """
        query = ""
        for a in acc_list:
            query += str(a) + " "
        base_dict = {
                "from": "ACC+ID",
                "to": "ACC",
                "query": query,
                "compress": "no",
                "force": "no",
                "sort": "score",
                "desc": "",
                "fil": "",
                "format": format
            }
        if format == "tab":
            base_dict["columns"] = "id,entry name,reviewed,protein names,genes,organism,length,database(RefSeq)," \
                                   "organism-id,go-id,go(cellular component),comment(SUBCELLULAR LOCATION)," \
                                   "feature(TOPOLOGICAL_DOMAIN),feature(GLYCOSYLATION),comment(MASS SPECTROMETRY)," \
                                   "sequence,feature(ALTERNATIVE SEQUENCE),comment(ALTERNATIVE PRODUCTS) "
        if include_isoform:
            base_dict["include"] = "yes"
        return base_dict

    def get(self, params):
        r = requests.get(self.base_url, params=params, headers=self.headers)
        return r

    def parse(self, format="fasta"):
        for i in range(0, self.total_input, 300):
            if (i + 300) <= self.total_input:

                params = self.create_params(self.acc_list[i: i + 300], format=format)
            else:

                params = self.create_params(self.acc_list[i: self.total_input], format=format)

            yield self.get(params).text


if __name__ == "__main__":
    import pandas as pd
    from io import StringIO
    msstats = pd.read_csv("out.csv_msstats.csv", index_col=0)

    for i, r in msstats.iterrows():
        seq = UniprotSequence(r["Protein"], True)
        msstats.at[i, "Accession"] = str(seq)
    accessions = msstats["Accession"].unique()
    parser = UniprotParser(accessions, True)

    data = []
    for i in parser.parse("tab"):
        frame = pd.read_csv(StringIO(i), sep="\t")
        frame = frame.rename(columns={frame.columns[-1]: "query"})
        data.append(frame)
    data = pd.concat(data, ignore_index=True)
    unmatched = []
    for a in accessions:
        if a not in data["query"].values:
            unmatched.append(a)
    if unmatched:
        print("Non-Uniprot ID found:", unmatched)
```
