## Supplementary material for "GlypNirO: An automated workflow for quantitative *N*- and *O*-linked glycoproteomic data analysis": glypniro-master.zip: main.py

```
import argparse
import pandas as pd
from glypnirO.common import GlypnirO

parser = argparse.ArgumentParser(description="Automated workflow for processing and combining Byonic and PD output")

parser.add_argument("-i",
                    "-input_file",
                    type=str,
                    help="Filepath to experiment description file where each row has 4 columns, condition_id, "
                         "replicate_id, filename, and area_filename")
parser.add_argument("-o",
                    "-output",
                    type=str,
                    help="Filepath to output")
parser.add_argument("-s",
                    "--score_cutoff",
                    type=float,
                    help="Filter the data by specific Byonic cut-off quality score",
                    default=200)

parser.add_argument("-t",
                    "--trust_byonic",
                    help="Enable site specific glycan position parsing from Byonic glycan assignment",
                    action="store_true")

parser.add_argument("-g",
                    "--get_uniprot",
                    help="Required internet connection. Enable parsing of UniProt ID from protein name and request "
                         "the original protein name from the UniProt databas.",
                    action="store_true")

if __name__ == "__main__":

    args = parser.parse_args()
    print("Input: " + args.i)
    print("Output: " + args.o)
    ex = GlypnirO(trust_byonic=args.trust_byonic, get_uniprot=args.get_uniprot)
    for i, r in ex.add_batch_component(args.i, args.score_cutoff):
        print(r)

    ex.process_components()
    result = ex.analyze_components()
    with pd.ExcelWriter(args.o) as writer:
        print("Writing Occupancy data to excel sheets.")
        result["Occupancy"].to_excel(writer, sheet_name="Unglyco_and_Glycoforms_Prop")
        result["Occupancy_Without_Proportion_U"].to_excel(writer, sheet_name="Unglyco_and_Glycoforms_Sep")
        result["Occupancy_With_U"].to_excel(writer, sheet_name="Unglycosylated")
        result["Glycoforms"].to_excel(writer, sheet_name="Glycoforms")
    print("Completed.")
```
