## Supplementary material for "GlypNirO: An automated workflow for quantitative *N*- and *O*-linked glycoproteomic data analysis": glypniro-master.zip: README.md

# GlypNirO

A automated script for processing and combining Byonic and PD standard output.

# Requirements

Pythons 3.4+ with the following packages installed.
- pandas
- requests
- xlrd
- openpyxl

# Usage

The program can be run from the location of the script using `python main.py`. The following parameters can be used for operating the script within the commandline.

|Paremeters|Descriptions|
|---------:|-----------:|
|`-i`, `-input_file`|Filepath to an xlsx file describing the experiment.|
|`-o`, `-output`|Filepath to the output xlsx file for the analysis.|
|`-s`, `--score_cutoff`|(Optional) Default=200. Cutoff score for filtering of Byonic output|
|`-t`, `--trust_byonic`|(Optional) Instruct the script to trust glycan position assignment and used them for area under the curve calculation.|
|`-g`, `--get_uniprot`|Using the `requests` module to access and parse protein name from uniprot accession ids of the proteins of those within the dataset.|

## Input file format

The input file used in the `-i` parameter should have the following format.

Ex:
|condition_id|replicate_id|filename|area_filename|
|---------:|-----------:|---------:|-----------:|
|Depleted_Plasma_HCC|1|\data\NoMutliFuc_Nlink_10_20ppm_05Da_Depleted_Plasma_HCC_1.raw_Byonic.xlsx|\data\Depleted_Plasma_HCC_1_MSnSpectrumInfo.txt|
|Depleted_Plasma_HCC|2|\data\NoMutliFuc_Nlink_10_20ppm_05Da_Depleted_Plasma_HCC_2.raw_Byonic.xlsx|\data\Depleted_Plasma_HCC_2_MSnSpectrumInfo.txt|
|Depleted_Plasma_HCC|3|\data\NoMutliFuc_Nlink_10_20ppm_05Da_Depleted_Plasma_HCC_3.raw_Byonic.xlsx|\data\Depleted_Plasma_HCC_3_MSnSpectrumInfo.txt|
|Depleted_Plasma_Nor|1|\data\NoMutliFuc_Nlink_10_20ppm_05Da_Depleted_Plasma_Nor_1.raw_Byonic.xlsx|\data\Depleted_Plasma_Nor_1_MSnSpectrumInfo.txt|	
|Depleted_Plasma_Nor|2|\data\NoMutliFuc_Nlink_10_20ppm_05Da_Depleted_Plasma_Nor_2.raw_Byonic.xlsx|\data\Depleted_Plasma_Nor_2_MSnSpectrumInfo.txt|	
|Depleted_Plasma_Nor|3|\data\NoMutliFuc_Nlink_10_20ppm_05Da_Depleted_Plasma_Nor_3.raw_Byonic.xlsx|\data\Depleted_Plasma_Nor_3_MSnSpectrumInfo.txt|	
|Non_Depleted_Plasma_HCC|1|\data\NoMutliFuc_Nlink_10_20ppm_05Da_Non_Depleted_Plasma_HCC_1.raw_Byonic.xlsx|\data\Non_Depleted_Plasma_HCC_1_MSnSpectrumInfo.txt|	
|Non_Depleted_Plasma_HCC|2|\data\NoMutliFuc_Nlink_10_20ppm_05Da_Non_Depleted_Plasma_HCC_2.raw_Byonic.xlsx|\data\Non_Depleted_Plasma_HCC_2_MSnSpectrumInfo.txt|
|Non_Depleted_Plasma_HCC|3|\data\NoMutliFuc_Nlink_10_20ppm_05Da_Non_Depleted_Plasma_HCC_3.raw_Byonic.xlsx|\data\Non_Depleted_Plasma_HCC_3_MSnSpectrumInfo.txt|	
|Non_Depleted_Plasma_Nor|1|\data\NoMutliFuc_Nlink_10_20ppm_05Da_Non_Depleted_Plasma_Nor_1.raw_Byonic.xlsx|\data\Non_Depleted_Plasma_Nor_1_MSnSpectrumInfo.txt|
|Non_Depleted_Plasma_Nor|2|\data\NoMutliFuc_Nlink_10_20ppm_05Da_Non_Depleted_Plasma_Nor_2.raw_Byonic.xlsx|\data\Non_Depleted_Plasma_Nor_2_MSnSpectrumInfo.txt|	
|Non_Depleted_Plasma_Nor|3|\data\NoMutliFuc_Nlink_10_20ppm_05Da_Non_Depleted_Plasma_Nor_3.raw_Byonic.xlsx|\data\Non_Depleted_Plasma_Nor_3_MSnSpectrumInfo.txt|

The rows within the input file should not have to follow any particular order however the columns have to contain the necessary content:
- `condition_id` condition label of the experiment
- `replicate_id` replicate label for the experiment
- `filename` filepath to the Byonic .xlsx output file of the experiment.
- `area_filename` filepath to the PD tabulated output file of the experiment.

# Output

The output of the script is a single .xlsx file with 4 sheets. 
- The first one is the output where we calculate the proportion by combining both the glycosylated and unglycosylated glycoform data.
- The second one is the output where the proportion of glycosylated was calculated without the unglycosylated data while the unglycosylated data was calculated similar to the first sheet.
- The third one is the only the filter of the glycosylated output from the second sheet.
- The forth one is the filter for only of the unglycosylated output from the first sheet.

# Example

`python main.py -i test_experiment.xlsx -o test_output.xlsx -t -g`

The above command would instruct the script to use the `test_experiment.xlsx` file as input file and output as `test_output.xlsx`. 
Inclusion of `-t` would mean that we trust Byonic assignment of glycan position and shall use them for calculation of that specific glycoform AUC within the proteins.
Inclusion of `-g` would instruct the script to connect to the UniProt online database and attempt to parse protein name from the UniProt accession id contain in the protein name within the Byonic file.

`python main.py -i test_experiment.xlsx -o test_output.xlsx`

The above command would instruct the script to use the `test_experiment.xlsx` file as input file and output as `test_output.xlsx`.
Without `-t` optional parameter, we only use the information of what glycans were found but not assigning them any positions. The AUC will only be combined for those PSMs with the same peptide sequence and glycan combination.
