## Supplementary material for "GlypNirO: An automated workflow for quantitative *N*- and *O*-linked glycoproteomic data analysis": glypniro-master.zip: amino_acid.py

```
from sequal.base_block import BaseBlock
from sequal.modification import Modification
from sequal.resources import AA_mass


# Basic amino acid block object. Can cary position, modifications and amino acid value.
class AminoAcid(BaseBlock):
    def __init__(self, value, position=None, mass=None):
        """

        :type mass: float
        mass of the amino acid if mass is not specified, it will try to infer the mass from the internal hard code mass dictionary of amino acid
        :type position: int
        position of amino acid residue that this block belong to
        :type value: str
        name of the amino acid residue for this block
        """
        super().__init__(value, position, branch=False, mass=mass)
        self.mods = []
        if not self.mass:
            if value in AA_mass:
                self.mass = AA_mass[value]

    # Adding modification to the mods list of this amino acid block
    def set_modification(self, i: Modification):
        self.mods.append(i)

    def __repr__(self):
        s = self.value
        for i in self.mods:
            s += "[{}]".format(i.value)
        return s

    def __str__(self):
        s = self.value
        for i in self.mods:
            s += "[{}]".format(i.value)
        return s
```
