## Supplementary material for "GlypNirO: An automated workflow for quantitative *N*- and *O*-linked glycoproteomic data analysis": glypniro-master.zip: base_block.py

```
# Base block object to be used for blocks that can have position and mass properties
class BaseBlock:
    def __init__(self, value, position, branch=False, mass=None):
        """
        :type mass: float
        mass of the block
        :type branch: bool
        whether or not this block is a branch of another block
        :type position: int
        position of the block within a chain
        :type value: str
        name of the block


        """
        self.value = value
        self.position = position
        self.branch = branch
        self.mass = mass
        self.extra = None

    def __str__(self):
        return self.value

    def __repr__(self):
        return self.value
```
