## Supplementary material for "GlypNirO: An automated workflow for quantitative *N*- and *O*-linked glycoproteomic data analysis": glypniro-master.zip: ion.py

```
from sequal.sequence import Sequence
from sequal.mass import calculate_mass
from sequal.resources import proton

modifier = {
    "b": -18-19,
}

# Ion fragment sequence object, similar property to the original sequence object
class Ion(Sequence):
    def __init__(self, seq, charge=1, ion_type=None, fragment_number=None):
        """

        :type fragment_number: int
        number of the transition
        :type ion_type: str
        name of transition type
        :type charge: int
        ion charge
        :type seq: Sequence
        Sequence object to be converted into an Ion fragment
        """
        super().__init__(seq)
        self.charge = charge
        self.ion_type = ion_type
        self.fragment_number = fragment_number
        self.mods = {}
        self.has_labile = False
        # Iterating through each amino acid position and build a modification list for the ion
        for i, aa in enumerate(self.seq):
            for m in aa.mods:
                if i not in self.mods:
                    self.mods[i] = []
                self.mods[i].append(m)
                if m.labile:
                    self.has_labile = True

    # Calculate the mz of the ion
    def mz_calculate(self, charge=None, with_water=False, extra_mass=0):
        """

        :type extra_mass: float
        extra modification of mass that is not represented within the sequence
        :type with_water: bool
        whether the mass will be calculated with or without water
        :type charge: int

        """
        if not charge:
            charge = self.charge
        m = calculate_mass(self.seq, with_water=with_water) + extra_mass

        # Charge is calculated with the hardcoded mass of protons
        mi = (m + charge*proton)/charge
        return mi
```
