## Supplementary material for "GlypNirO: An automated workflow for quantitative *N*- and *O*-linked glycoproteomic data analysis": glypniro-master.zip: mass.py

```
from sequal import resources

# Calculate the mass of a Sequence object using a mass dictionary of
def calculate_mass(seq, mass_dict=None, N_terminus=0, O_terminus=0, with_water=True):
    """

    :rtype: float
    mass of the sequence calculated using the input parameters
    :type with_water: bool
    whether or not to add the mass of water
    :type O_terminus: int or float
    mass at the C-terminus of the sequence
    :type N_terminus: int or float
    mass at the N-terminus of the sequence
    :type mass_dict: dict
    a dictionary of mass containing mass of potential modifications and amino acids
    :type seq: sequal.sequence.Sequence
    a Sequence object
    """
    mass = 0
    if with_water:
        mass += resources.H*2 + resources.O
    for i in seq:
        if not i.mass:
            if mass_dict:
                if i.value in mass_dict:
                    mass += mass_dict[i.value]
                else:
                    raise ValueError('Block {} not found in mass_dict'.format(i.value))
            else:
                raise ValueError('Block {} mass is not available in mass attribute and no additional mass_dict was supplied'.format(i.value))
        else:
            mass += i.mass
        if i.mods:
            for m in i.mods:
                if m.mass != 0 and not m.mass:
                    if mass_dict:
                        if m.value in mass_dict:
                            mass += mass_dict[m.value]
                        else:
                            raise ValueError('Block {} not found in mass_dict'.format(m.value))
                    else:
                        raise ValueError(
                            'Block {} mass is not available in mass attribute and no additional mass_dict was supplied'.format(
                                m.value))
                else:
                    mass += m.mass
    return mass + N_terminus + O_terminus
```
