## Supplementary material for "GlypNirO: An automated workflow for quantitative *N*- and *O*-linked glycoproteomic data analysis": glypniro-master.zip: mass_spectrometry.py

```
from sequal.ion import Ion

ax = "ax"
by = "by"
cz = "cz"

# calculate non-labile modifications and yield associated transition
# For example "by" would yield a tuple of "b" and "y" transitions.
def fragment_non_labile(sequence, fragment_type):
    for i in range(1, sequence.seq_length, 1):
        left = Ion(sequence[:i], fragment_number=i, ion_type=fragment_type[0])
        right = Ion(sequence[i:], fragment_number=sequence.seq_length-i, ion_type=fragment_type[1])
        yield left, right

# calculate all labile modification variants for the sequence and its associated labile modifications
def fragment_labile(sequence):
    fragment_number = 0
    for p in sequence.mods:
        for i in sequence.mods[p]:
            if i.labile:
                fragment_number += i.labile_number
    return Ion(sequence, fragment_number=fragment_number, ion_type="Y")


class FragmentFactory:
    def __init__(self, fragment_type, ignore=None):
        self.fragment_type = fragment_type
        if ignore:
            self.ignore = ignore
        else:
            self.ignore = []

    def set_ignore(self, ignore):
        self.ignore = ignore
```
