## Supplementary material for "GlypNirO: An automated workflow for quantitative *N*- and *O*-linked glycoproteomic data analysis": glypniro-master.zip: mass_spectrometry_test.py

```
import unittest

from sequal.mass_spectrometry import fragment_non_labile, fragment_labile
from sequal.modification import Modification
from sequal.sequence import ModdedSequenceGenerator, Sequence

nsequon = Modification("HexNAc",regex_pattern="N[^P][S|T]", mod_type="variable", labile=True, labile_number=1, mass=203)
osequon = Modification("Mannose",regex_pattern="[S|T]", mod_type="variable", labile=True)
sulfation = Modification("Sulfation",regex_pattern="S", mod_type="variable", labile=True)
carbox = Modification("Carboxylation",regex_pattern="E", mod_type="variable", labile=True, labile_number=1)
carbox2 = Modification("Carboxylation2", regex_pattern="E", mod_type="variable", labile=True, mass=43.98983)
propiona = Modification("Propionamide", regex_pattern="C", mod_type="static", mass=71)


class MassSpecTestCase(unittest.TestCase):
    def test_fragment_non_labile(self):
        seq = "TECSNTT"
        static_mods = [propiona]
        variable_mods = [nsequon]
        g = ModdedSequenceGenerator(seq, variable_mods, static_mods)
        for i in g.generate():
            print(i)
            s = Sequence(seq, mods=i)
            print(s)
            for b, y in fragment_non_labile(s, "by"):
                print(b, "b{}".format(b.fragment_number))
                print(y, "y{}".format(y.fragment_number))

    def test_fragment_labile(self):
        seq = "TECSNTT"
        static_mods = [propiona]
        variable_mods = [nsequon]
        g = ModdedSequenceGenerator(seq, variable_mods, static_mods)
        for i in g.generate():
            s = Sequence(seq, mods=i)
            ion = fragment_labile(s)
            if ion.has_labile:
                print(ion, "Y{}".format(ion.fragment_number))
                print(ion.mz_calculate(1))


if __name__ == '__main__':
    unittest.main()
```
