## Supplementary material for "GlypNirO: An automated workflow for quantitative *N*- and *O*-linked glycoproteomic data analysis": glypniro-master.zip: mass_test.py

```
import unittest

from sequal.mass import calculate_mass
from sequal.mass_spectrometry import fragment_non_labile
from sequal.modification import Modification
from sequal.sequence import ModdedSequenceGenerator, Sequence

nsequon = Modification("HexNAc", regex_pattern="N[^P][S|T]", mod_type="variable", labile=True, mass=203.0794)
osequon = Modification("Mannose", regex_pattern="[S|T]", mod_type="variable", labile=True, mass=162.05)
sulfation = Modification("Sulfation", regex_pattern="S", mod_type="variable", labile=True, mass=79.95680)
carbox = Modification("Carboxylation", regex_pattern="E", mod_type="variable", labile=True, mass=43.98983)
propiona = Modification("Propionamide", regex_pattern="C", mod_type="static", mass=71.037114)


class MassTestCase(unittest.TestCase):
    def test_mass_calculation(self):
        seq = "TECSNTT"
        static_mods = [propiona]
        variable_mods = [nsequon]
        g = ModdedSequenceGenerator(seq, variable_mods, static_mods)
        for i in g.generate():
            seqs = Sequence(seq, mods=i)
            print(seqs)
            mass = calculate_mass(seqs, N_terminus=0, O_terminus=0)
            print(mass)

    def test_fragment_mass_calculation(self):
        seq = "TECSNTT"
        static_mods = [propiona]
        variable_mods = [nsequon]
        g = ModdedSequenceGenerator(seq, variable_mods, static_mods)
        for i in g.generate():
            seqs = Sequence(seq, mods=i)
            for b, y in fragment_non_labile(seqs, "by"):
                print(b, "b{}".format(b.fragment_number))
                mass_b = calculate_mass(b, N_terminus=1.007825, O_terminus=0, with_water=False)
                print(mass_b)
                print(y, "y{}".format(y.fragment_number))
                mass_y = calculate_mass(y, N_terminus=0, O_terminus=15.99491463+1.007825, with_water=False)
                print(mass_y)

if __name__ == '__main__':
    unittest.main()
```
