## Supplementary material for "GlypNirO: An automated workflow for quantitative *N*- and *O*-linked glycoproteomic data analysis": glypniro-master.zip: modification.py

```
import re

from sequal.base_block import BaseBlock


# Base modification block.
class Modification(BaseBlock):
    def __init__(self, value, position=None, regex_pattern=None, full_name=None, mod_type="static", labile=False, labile_number=0, mass=0, all_filled=False):
        """

        :type all_filled: bool
        whether or not the modification could be always found at all the expected sites
        :type mass: float
        mass of the modification
        :type labile_number: int
        what is the order of the fragment in a labile fragmentation event
        :type labile: bool
        whether or not the modification is labile. Important for mass spectrometry fragmentation.
        :type mod_type: str
        modification type. Either "static" or "variable".
        :type full_name: str
        full name of modification
        :type regex_pattern: str
        regular expression pattern that can be used to represent where the modication is found at
        :type value: str
        short name of modification
        :type position: int
        Position of the modification on the block it belongs to. Should be int and not None if it is assigned to a
        block. None when using as unassigned.
        """
        super().__init__(value, position=position, branch=True, mass=mass)
        if regex_pattern:
            self.regex = re.compile(regex_pattern)
        else:
            self.regex = None
        if mod_type in {"static", "variable"}:
            self.mod_type = mod_type
        else:
            print("Type can only be 'static' or 'variable'")
            raise ValueError

        assert(type(labile) == bool)
        self.labile = labile
        self.labile_number = labile_number
        self.full_name = full_name
        self.all_fill = all_filled

    def __repr__(self):
        if not self.labile:
            return self.value
        else:
            return self.value + str(self.labile_number)

    def __str__(self):
        if not self.labile:
            return self.value
        else:
            return self.value + str(self.labile_number)

    def find_positions(self, seq):
        for i in self.regex.finditer(seq):
            res = len(i.groups())
            if res > 0:
                for r in range(0, res + 1):
                    yield i.start(r), i.end(r)
            else:
                yield i.start(), i.end()


# Internal used object that carry dictionary of modification, modification position and their name for quick reference
class ModificationMap:
    def __init__(self, seq, mods, ignore_positions=None, parse_position=True, mod_position_dict=None):
        self.ignore_positions = ignore_positions
        self.seq = seq
        self.mod_dict_by_name = {}
        if mod_position_dict:
            self.mod_position_dict = mod_position_dict

        for m in mods:
            self.mod_dict_by_name[str(m)] = m
            if parse_position:
                d = []
                for p_start, p_end in m.find_positions(self.seq):
                    if ignore_positions:
                        if p_start not in ignore_positions:
                            d.append(p_start)
                    else:
                        d.append(p_start)
                self.mod_position_dict[str(m)] = d
        # print(self.mod_position_dict)

    def get_mod_positions(self, mod_name):
        if mod_name in self.mod_position_dict:
            return self.mod_position_dict[mod_name]
        else:
            return None

    def get_mod(self, mod_name):
        if mod_name in self.mod_dict_by_name:
            return self.mod_dict_by_name[mod_name]
        else:
            return None
```
