## Supplementary material for "GlypNirO: An automated workflow for quantitative *N*- and *O*-linked glycoproteomic data analysis": glypniro-master.zip: modification_test.py

```
import unittest
from sequal.modification import Modification


class ModificationTestCase(unittest.TestCase):
    def test_find_positions(self):
        mod = Modification("HexNAc", regex_pattern="N[^P][S|T]")
        for ps, pe in mod.find_positions("TESNEST"):
            print(ps, pe)
            self.assertEqual(ps, 3, "HexNAc is at expected index position {}".format(ps))

class ModificationMapTestCase(unittest.TestCase):
    def test_map_creation(self):
        pass


if __name__ == '__main__':
    unittest.main()
```
