## Supplementary material for "GlypNirO: An automated workflow for quantitative *N*- and *O*-linked glycoproteomic data analysis": glypniro-master.zip: resources.py

```
# Hardcoded resources for use of calculating mass for sequences, mass spectrometry ion fragments and glycans

proton = 1.007277
H = 1.007825
O = 15.99491463
AA_mass = {"A": 71.037114,
           "R":	156.101111,
           "N": 114.042927,
           "D": 115.026943,
           "C": 103.009185,
           "E": 129.042593,
           "Q": 128.058578,
           "G": 57.021464,
           "H": 137.058912,
           "I": 113.084064,
           "L": 113.084064,
           "K": 128.094963,
           "M": 131.040485,
           "F": 147.068414,
           "P": 97.052764,
           "S": 87.032028,
           "T": 101.047679,
           "U": 150.95363,
           "W": 186.079313,
           "Y": 163.06332,
           "V": 99.068414}

glycan_block_dict = {"HexNAc": 203.079372520,
          "Hex": 162.0528234185,
          "Fuc": 146.057908799,
          "NeuAc": 291.0954165066,
          "Sulfo": 79.9568148602,
          "Phospho": 79.9663305228,
          "Pent": 132.0422587348,
          "NeuGc": 307.0903311261}
```
