## Supplementary material for "GlypNirO: An automated workflow for quantitative *N*- and *O*-linked glycoproteomic data analysis": glypniro-master.zip: sequence.py

```
import re
from typing import Set, Any, List

from sequal.amino_acid import AminoAcid
from sequal.modification import Modification, ModificationMap
from copy import deepcopy
import itertools
from json import dumps
mod_pattern = re.compile(r"[\(|\[]+([^\)]+)[\)|\]]+")
mod_enclosure_start = {"(", "[", "{"}
mod_enclosure_end = {")", "]", "}"}

# Base sequence object for peptide or protein sequences and their fragments
class Sequence:
    seq: List[Any]

    def __init__(self, seq, encoder=AminoAcid, mods=None, parse=True, parser_ignore=None, mod_position="right"):
        """
        :param mod_position
        Indicate the position of the modifications relative to the base block it is supposed to modify
        :type mod_position: str
        :param mods
        Dictionary whose keys are the positions within the sequence and values are array of modifications at those
        positions
        :type mods: dict
        :param encoder
        Class for encoding of sequence.
        :type encoder: BaseBlock
        :param seq
        String or array of strings or array of AminoAcid objects. The parser will recursively look over each string at
        deepest level and identify individual modifications or amino acids for processing
        :type seq: iterable
        Python iterable where the deepest level is a string
            
        """
        if type(seq) is not Sequence:
            if not mods:
                self.mods = {}
            else:
                self.mods = mods
            self.encoder = encoder
            if not parser_ignore:
                self.parser_ignore = []
            else:
                self.parser_ignore = parser_ignore
            self.seq = []
            current_mod = []
            current_position = 0
            if parse:
                self.sequence_parse(current_mod, current_position, mod_position, mods, seq)

        else:
            for k in seq.__dict__:
                if k != "mods":
                    setattr(self, k, deepcopy(seq.__dict__[k]))
        self.seq_length = len(self.seq)

    def __getitem__(self, key):
        return self.seq[key]

    def __len__(self):
        return self.seq_length

    def __repr__(self):
        a = ""
        for i in self.seq:
            a += str(i)
        return a

    def __str__(self):
        a = ""
        for i in self.seq:
            a += str(i)
        return a

    def sequence_parse(self, current_mod, current_position, mod_position, mods, seq):
        """
        :param seq: sequence input
        :param mods: external modification input
        :param mod_position: modification position relative to the modified residue
        :param current_position: current iterating amino acid position from the input sequence
        :type current_mod: List[Modification]
        """
        for b, m in self.__load_sequence_iter(iter(seq)):
            if not m:
                if mod_position == "left":
                    if type(b) == AminoAcid:
                        current_unit = b
                        current_unit.position = current_position
                    else:
                        current_unit = self.encoder(b, current_position)

                    if current_mod and not mods:
                        for i in current_mod:
                            current_unit.set_modification(i)
                    elif current_position in self.mods and current_unit:
                        if type(self.mods[current_position]) == Modification:
                            current_unit.set_modification(self.mods[current_position])
                        else:
                            for mod in self.mods[current_position]:
                                current_unit.set_modification(mod)

                    self.seq.append(deepcopy(current_unit))

                    current_mod = []
                if mod_position == "right":

                    if current_mod and not mods:
                        for i in current_mod:
                            self.seq[current_position - 1].set_modification(i)
                    if type(b) == AminoAcid:
                        current_unit = b
                        current_unit.position = current_position
                    else:
                        current_unit = self.encoder(b, current_position)

                    if current_position in self.mods and current_unit:
                        if type(self.mods[current_position]) == Modification:
                            current_unit.set_modification(self.mods[current_position])

                        else:
                            for mod in self.mods[current_position]:
                                current_unit.set_modification(mod)

                    self.seq.append(deepcopy(current_unit))

                    current_mod = []
                current_position += 1
            else:
                if not mods:
                    # current_mod.append(Modification(b[1:-1]))
                    if mod_position == "right":
                        self.seq[current_position-1].set_modification(Modification(b[1:-1]))
                    else:
                        current_mod.append(Modification(b[1:-1]))

    def __load_sequence_iter(self, seq=None, iter_seq=None):
        mod_open = 0
        block = ""
        mod = False
        if not iter_seq:
            iter_seq = iter(seq)
        for i in iter_seq:
            if type(i) == str:
                if i in mod_enclosure_start:
                    mod = True
                    mod_open += 1
                elif i in mod_enclosure_end:
                    mod_open -= 1
                block += i
            elif type(i) == AminoAcid:
                block = i
            else:
                yield from self.__load_sequence_iter(iter_seq=iter_seq)
            if mod_open == 0:
                yield (block, mod)
                mod = False
                block = ""

    def __iter__(self):
        self.current_iter_count = 0
        return self

    def __next__(self):
        if self.current_iter_count == self.seq_length:
            raise StopIteration
        result = self.seq[self.current_iter_count]
        self.current_iter_count += 1
        return result

    def add_modifications(self, mod_dict):
        for aa in self.seq:
            if aa.position in mod_dict:
                for mod in mod_dict[aa.position]:
                    aa.set_modification(mod)

    def to_stripped_string(self):
        """
        Return string of the sequence without any modification annotation
        :return: str
        """
        seq = ""
        for i in self.seq:
            seq += i.value
        return seq

    def to_string_customize(self, data, annotation_placement="right", block_separator="", annotation_enclose_characters=("[", "]"),
                            individual_annotation_enclose=False, individual_annotation_enclose_characters=("[", "]"),
                            individual_annotation_separator=""):
        """

        :rtype: str
        :param data: a dictionary where the key is the index position of the amino acid residue and the value is a
        iterable where containing the item needed to be included into the sequence.
        :param annotation_placement: whether the information should be included on the right of the left of the residue
        :param block_separator: separator between each block of annotation information to be included
        :param annotation_enclose_characters: enclosure characters for each annotation cluster
        :param individual_annotation_enclose: whether or not each individual annotation should be enclosed
        :param individual_annotation_enclose_characters: enclosure characters for each individual annotation
        :param individual_annotation_separator: separator for each individual annotation
        :return:
        """
        assert annotation_placement in {"left", "right"}
        seq = []
        for i in range(len(self.seq)):
            seq.append(self.seq[i].value)
            if i in data:
                annotation = []
                if individual_annotation_enclose:
                    for v in data[i]:
                        annotation.append("{}{}{}".format(individual_annotation_enclose_characters[0], v, individual_annotation_enclose_characters[1]))
                else:
                    annotation = data[i]
                if type(annotation) == str:
                    ann = annotation
                else:
                    ann = individual_annotation_separator.join(annotation)
                if annotation_enclose_characters:
                    seq.append("{}{}{}".format(annotation_enclose_characters[0], ann, annotation_enclose_characters[1]))
                else:
                    seq.append(individual_annotation_separator.join(ann))
        return block_separator.join(seq)

    def find_with_regex(self, motif, ignore=None):
        pattern = re.compile(motif)
        new_str = ""
        if ignore is not None:
            for i in range(len(ignore)):
                if not ignore[i]:
                    new_str += self.seq[i].value
        else:
            new_str = self.to_stripped_string()

        for i in pattern.finditer(new_str):

            if not i.groups():
                yield slice(i.start(), i.end())
            else:
                for m in range(1, len(i.groups()) + 1):
                    yield slice(i.start(m), i.end(m))

    def gaps(self):
        s = [False for i in range(len(self.seq))]
        for i in range(len(s)):
            if self.seq[i].value == '-':
                s[i] = True

        return s

    def count(self, char, start, end):
        return self.to_stripped_string().count(char, start, end)

def count_unique_elements(seq):
    elements = {}
    for i in seq:
        if i.value not in elements:
            elements[i.value] = 0
        elements[i.value] += 1
        if i.mods:
            for m in i.mods:
                if m.value not in elements:
                    elements[m.value] = 0
                elements[m.value] += 1
    return elements


def variable_position_placement_generator(positions):
    """
    Use itertools.product to generate a list of tuple with different number of 0 and 1. The length of the tuple is the
    length of the input positions.
    Using itertools.compress, for each output from itertools.product pairing with input positions, we generate a list of
    positions where only those with the same index as 1 would be yielded.

    :param positions: list of all identified positions for the modification on the sequence
    """
    for i in itertools.product([0, 1], repeat=len(positions)):
        yield list(itertools.compress(positions, i))


def ordered_serialize_position_dict(positions):
    return dumps(positions, sort_keys=True, default=str)


class ModdedSequenceGenerator:
    used_scenarios_set: Set[str]

    def __init__(self, seq, variable_mods=None, static_mods=None, used_scenarios=None, parse_mod_position=True, mod_position_dict=None, ignore_position=None):
        """
        Generator for creating modified sequences.
        :type used_scenarios: set
        :type static_mods: List[Modification]
        :type variable_mods: List[Modification]
        :type seq: str
        """
        self.seq = seq
        if static_mods:
            self.static_mods = static_mods

            self.static_map = ModificationMap(seq, static_mods, parse_position=parse_mod_position, mod_position_dict=mod_position_dict)
            self.static_mod_position_dict = self.static_mod_generate()
        else:
            self.static_mod_position_dict = {}
        if ignore_position:
            self.ignore_position = ignore_position
        else:
            self.ignore_position = set()

        for i in self.static_mod_position_dict:
            self.ignore_position.add(i)

        if variable_mods:
            self.variable_mods = variable_mods
            if self.static_mod_position_dict:
                self.variable_map = ModificationMap(seq, variable_mods, ignore_positions=self.ignore_position, parse_position=parse_mod_position, mod_position_dict=mod_position_dict)
            else:
                self.variable_map = ModificationMap(seq, variable_mods)
            self.variable_mod_number = len(variable_mods)
        else:
            self.variable_mods = None

        self.variable_map_scenarios = {}
        if used_scenarios:
            self.used_scenarios_set = used_scenarios
        else:
            self.used_scenarios_set = set()

    def generate(self):
        if self.variable_mods:
            self.variable_mod_generate_scenarios()
            for i in self.explore_scenarios():
                a = dict(self.static_mod_position_dict)
                a.update(i)
                serialized_a = ordered_serialize_position_dict(a)
                if serialized_a not in self.used_scenarios_set:
                    self.used_scenarios_set.add(serialized_a)
                    yield a
        else:
            serialized_a = ordered_serialize_position_dict(self.static_mod_position_dict)
            if serialized_a not in self.used_scenarios_set:
                yield self.static_mod_position_dict

    def static_mod_generate(self):
        position_dict = {}

        for m in self.static_mods:

            for pm in self.static_map.get_mod_positions(str(m)):
                if pm not in position_dict:
                    position_dict[pm] = []
                position_dict[pm].append(m)
        return position_dict

    def variable_mod_generate_scenarios(self):
        """
        Recursively generating all possible position compositions for each variable modification and add them to
        self.variable_map_scenarios dictionary where key is the value attr of the modification while the value is the
        position list
        """
        for i in self.variable_mods:
            positions = self.variable_map.get_mod_positions(str(i))
            if i.value not in self.variable_map_scenarios:
                if not i.all_fill:
                    self.variable_map_scenarios[i.value] = list(
                        variable_position_placement_generator(positions))
                else:
                    self.variable_map_scenarios[i.value] = [[], positions]


    def explore_scenarios(self, current_mod=0, mod=None):
        if mod is None:
            mod = {}
        for pos in self.variable_map_scenarios[self.variable_mods[current_mod].value]:
            temp_dict = deepcopy(mod)
            if pos:
                for p in pos:
                    if p not in temp_dict:
                        temp_dict[p] = [self.variable_mods[current_mod]]
                    if current_mod != self.variable_mod_number - 1:
                        yield from self.explore_scenarios(current_mod + 1, temp_dict)
                    else:
                        yield temp_dict
            else:
                if current_mod != self.variable_mod_number - 1:
                    yield from self.explore_scenarios(current_mod + 1, temp_dict)
                else:
                    yield temp_dict
```
