## Supplementary material for "GlypNirO: An automated workflow for quantitative *N*- and *O*-linked glycoproteomic data analysis": glypniro-master.zip: sequence_test.py

```
import unittest

from sequal.modification import Modification
from sequal.sequence import Sequence, ModdedSequenceGenerator

nsequon = Modification("HexNAc",regex_pattern="N[^P][S|T]", mod_type="variable", labile=True)
osequon = Modification("Mannose",regex_pattern="[S|T]", mod_type="variable", labile=True)
sulfation = Modification("Sulfation",regex_pattern="S", mod_type="variable", labile=True)
carbox = Modification("Carboxylation",regex_pattern="E", mod_type="variable", labile=True)
carbox2 = Modification("Carboxylation2", regex_pattern="E", mod_type="variable", labile=True, mass=43.98983)
propiona = Modification("Propionamide", regex_pattern="C", mod_type="static")


class TestAASequence(unittest.TestCase):
    def test_normal_sequence(self):
        seq = Sequence("TESTEST")
        print(seq.seq)
        print(seq[0:2])

    def test_mod_rightseq(self):
        seq = Sequence("TEN[HexNAc]ST")
        for i in seq.seq:
            print(i, i.mods)

    def test_two_mod_rightseq(self):
        seq = Sequence("TEN[HexNAc][HexNAc]ST")
        for i in seq.seq:
            print(i, i.mods)

    def test_mod_leftseq(self):
        seq = Sequence("TE[HexNAc]NST", mod_position="left")
        for i in seq.seq:
            print(i, i.mods)

    def test_two_mod_leftseq(self):
        seq = Sequence("TE[HexNAc][HexNAc]NST", mod_position="left")
        for i in seq.seq:
            print(i, i.mods)

    def test_custom_string(self):
        seq = Sequence("TENST")
        a = {1:"tes", 2:["1", "200"]}
        print(seq.to_string_customize(a, individual_annotation_enclose=False, individual_annotation_separator="."))


class TestModdedSequence(unittest.TestCase):
    def test_variable_mod_generator(self):
        seq = "TESNSTT"
        mods = [nsequon, osequon, carbox]
        g = ModdedSequenceGenerator(seq, mods, [])
        print(g.variable_map.mod_position_dict)
        for i in g.generate():
            print(i)

    def test_static_mod_generator(self):
        seq = "TECSNTT"
        mods = [propiona]
        g = ModdedSequenceGenerator(seq, static_mods=mods)
        for i in g.generate():
            print(i)

    def test_static_and_variable_mod_generator(self):
        seq = "TECSNTT"
        static_mods = [propiona]
        variable_mods = [nsequon, osequon, carbox, carbox2]
        g = ModdedSequenceGenerator(seq, variable_mods, static_mods)
        for i in g.generate():
            print(i)

if __name__ == '__main__':
    unittest.main()
```
